# Profiling sorghum-microbe interactions with a specialized photoaffinity probe identifies key sorgoleone binders in *Acinetobacter pittii*

**DOI:** 10.1101/2023.05.31.542302

**Authors:** Jared O. Kroll, Elise M. Van Fossen, Lindsey N. Anderson, Andrew D. McNaughton, Daisy Herrera, Yasuhiro Oda, Andrew J. Wilson, William C. Nelson, Neeraj Kumar, Andrew R. Frank, Joshua R. Elmore, Pubudu Handakumbura, Vivian S. Lin, Robert G. Egbert

**Affiliations:** Energy Processes and Materials Division, Pacific Northwest National Laboratory, Richland, Washington, USA; Biological Sciences Division, Pacific Northwest National Laboratory, Richland, Washington, USA; Environmental and Molecular Sciences Division, Pacific Northwest National Laboratory, Richland, Washington, USA; Department of Microbiology, University of Washington, Seattle, Washington, USA

## Abstract

Sorghum (*Sorghum bicolor*) is a major food and bioenergy grass species cultivated worldwide. To promote more robust and sustainable growth of this important crop, we need a deeper understanding of the plant-microbe interactions between sorghum and soil microbial communities that benefit plant host resiliency and enhance nutrient acquisition. The release of specific metabolites from plant roots, or root exudation, drives these plant-microbe interactions, but the molecular pathways by which root exudates shape the sorghum rhizosphere microbiome require further elucidation. To investigate these complex interkingdom interactions in the sorghum rhizosphere, we developed a photoaffinity probe based on sorgoleone, a hydrophobic secondary metabolite and allelochemical produced in sorghum seedling root exudates. Here, we apply a new synthetic sorgoleone diazirine alkyne photoaffinity probe (SoDA-PAL) to the identification of sorgoleone-binding proteins in *Acinetobacter pittii* SO1, a potential plant growth promoting microbe derived from Sorghum bicolor rhizosphere soil. Competitive photoaffinity labeling of *A. pittii* whole cell lysates with SoDA-PAL identified 137 statistically enriched proteins that were complementary to a previously identified gene cluster involved in sorgoleone catabolism. Proteins identified by SoDA-PAL included a select set of putative transporters, transcription regulators, and a subset of proteins with lipid and secondary metabolic activities. We confirm binding of SoDA-PAL to a putative hydrolase in the α/β fold family (OH685_09420) through structural bioinformatics and in-vitro recombinant protein analysis. This photoaffinity labeling approach using metabolite-based probes can be extended in the future to proteomic profiling of complex rhizosphere microbiomes to discover genes that can be leveraged to promote beneficial plant-microbe interactions.

**Importance:** Here we demonstrate a photoaffinity-based chemical probe modeled after sorgoleone, a known secondary metabolite released from the roots of sorghum, can be used to dissect complicated plant-microbe interactions. Applying this probe to the sorghum-associated bacterium *Acinetobacter pittii* identified diverse proteins that directly interact with sorgoleone. We show that probe labeling is dose-dependent and is sensitive to competition with purified sorgoleone, demonstrating the probe is selective for protein targets that directly interact with sorgoleone. By using the probe to broadly profile proteins that interact with sorgoleone, we identified bacterial catabolic pathways, unintuitive transcriptional regulation pathways, and vital exchange mechanisms involving transporters that may be involved in sorgoleone utilization and cellular response toward this plant metabolite. We envision that this workflow will expand our understanding of the sorghum root exudate interactome and elucidate the molecular mechanisms by which specific metabolites shape the sorghum rhizosphere microbiome.

## Introduction

Sorghum (*Sorghum bicolor*) is a C_4_ grass species grown worldwide for its grain, making it the 5^th^ most important cereal crop. This widely cultivated grass can be productive under drought and low nutrient input conditions, making it an appealing species for bioenergy production.^1, 2^ Sorghum produces and exudes sorgoleone—a hydrophobic secondary metabolite and allelochemical—from its roots in significant quantities.^3, 4^ Sorgoleone can dramatically reduce growth of other proximal plant species by inhibiting electron transfer reaction photosystem II and other cellular functions.^4–6^ Sorgoleone production is particularly high during early sorghum growth stages, providing seedlings with a competitive advantage for limited space and nutrients.^7^ Given the high abundance of this carbon-rich phytochemical in sorghum root exudates, we hypothesized that specialized bacteria in the sorghum rhizosphere may be adapted to utilize sorgoleone as a carbon source for growth. Previous studies have observed that sorgoleone persists to some extent in soil, particularly within short time periods (1 hr.), but it is significantly degraded in soils over longer incubation periods.^8, 9^ This suggests that members of certain soil communities can metabolize this lipophilic constituent of sorghum root exudates, though the products of sorgoleone degradation in the sorghum rhizosphere have not yet been identified.^4^ Plant root exudation plays a critical role in shaping rhizosphere microbiomes,^10, 11^ and recently published reports have characterized the impact of sorgoleone on soil microbial communities.^12–14^ Understanding the catabolic pathways involved in sorgoleone utilization will provide deeper insight into the metabolic interactions between sorghum and its root-associated microbiome, which supports strategies to identify and engineer plant growth promoting bacteria (PGPB) for sorghum.

While a variety of proteins have been identified as molecular targets of sorgoleone and related benzoquinones in plant systems,^15–17^ only a few recent studies have examined the interaction of sorgoleone-utilizing microbial species with sorgoleone.^12^ Sorgoleone reduces rates of nitrification by influencing populations of nitrifying bacteria in the soil microbial community through the inhibition of key enzymes such as ammonia monooxygenase and hydroxylamine oxidoreductase.^13, 14^ While some studies have focused on known interactions between benzoquinones and specific proteins, as well as sorgoleone-induced changes at the microbial community level,^12^ the metabolic pathways involving catabolism of sorgoleone remain largely unknown. Recently, our group has identified specific genes response for utilization of sorgoleone as a sole carbon source in bacterial isolates from the sorghum rhizosphere and specifically in *Acinetobacter pittii* strain SO1 (TaxonID:48296).^18^ *Acinetobacter* have been shown to be beneficial for plant growth by engaging in beneficial nutrient cycling and by protecting against pathogens and abiotic stress.^19, 20^

These previous studies demonstrate that sorgoleone can have disparate impacts on the growth and functions of different microbial species, highlighting the potential of sorgoleone as a promising modulator of microbial functions and interactions in the sorghum rhizosphere. However, unraveling rhizosphere microbiome interactions with specific small molecule components of root exudates is a challenging endeavor due to the complexity of soil and plant-associated microbial communities. The selection of specific microbes that can grow on a specific compound can be achieved through sequential culturing on minimal media with the compound of interest as the sole carbon source.^21–23^ This strategy has helped to advance our understanding of microbes that can effectively utilize those compounds and develop model consortia for laboratory studies,^24^ but such approaches may not be able to identify species that do not grow well when isolated from their native environment.

The field of chemical biology provides new avenues to investigate plant-microbe interactions through the development and application of tailored chemical probes.^25^ Three distinct components are required to generate a functional chemical probe for identifying protein binders for plant metabolites: a recognition group for binding to specific proteins of interest, a reactive group for covalently labeling the protein when bound to the probe, and a reporter group for providing a readout of the labeling event.^26–28^ In lieu of a reporter group, an azide or alkyne “click” tag can be used for later attachment of various reporters using click chemistry.^29–31^ These small, bio-orthogonal tags allow for a fluorophore to be attached for fluorescence imaging of protein bands resolved by gel electrophoresis or for a biotin to be attached for streptavidin enrichment and subsequent digestion for bottom-up proteomics analysis. The flexibility imparted by using “click” tags in chemical probe designs make these an appealing strategy for multimodal analysis of probe-labeled protein targets in downstream steps of the workflow.^32, 33^ Due to their modular nature, chemical probes can be tailored for diverse applications and metabolites of interest. Furthermore, since chemical probes can be directly applied to biological systems without requiring genetic modification, they are well-suited for profiling complex systems, such as microbial communities, and can be used to enrich a subset of the proteome or a functionally active population for increased sensitivity and selectivity for low abundance analytes or community members.^29–31^

To this end, we designed and synthesized a photoaffinity probe based on the structure of sorgoleone to identify the proteins that may be involved in sorgoleone catabolism and enable a more global search for microbes that interact with sorgoleone **(Figure 1).** Photoaffinity probes are chemical probes that employ a photo-crosslinking reactive group to covalently attach the probe to its protein targets.^34–36^ After treating a complex proteome or sample with the photoaffinity probe, the sample is irradiated with light, converting the probe’s photo-crosslinking group to a highly reactive intermediate that can insert into nearby molecules. This photoaffinity labeling (PAL) approach has been used to study a wide array of different protein-ligand interactions and is particularly useful for discovery of previously unknown protein-small molecule interactions.^37–40^ By reporting on the specific physical interactions between proteins and metabolite ligands, PAL enables enrichment of even low abundance proteins, providing valuable protein-ligand interaction information beyond global protein abundance measurements alone. Previous studies using PAL probes based on plant-derived small molecules such as hormones and metabolites have specifically focused on identifying plant proteins that interact with these compounds, such as stevia,^41^ cytokinins,^42^ salicylic acid,^43^ indole-3-acetic acid,^44^ and gibberellin.^45^ Few studies have used photoaffinity probes to investigate plant-microbe interactions, in part due to the challenges in generating chemical probes based on plant-derived metabolites. Herein, we report the application of a sorgoleone diazirine alkyne photoaffinity probe (SoDA-PAL) based on the hydrophobic plant secondary metabolite sorgoleone for profiling proteins associated with sorgoleone utilization in a sorghum rhizosphere associated bacterium and demonstrate the utility of this probe-based approach for investigation of plant-microbe interactions.

**Figure 1.**
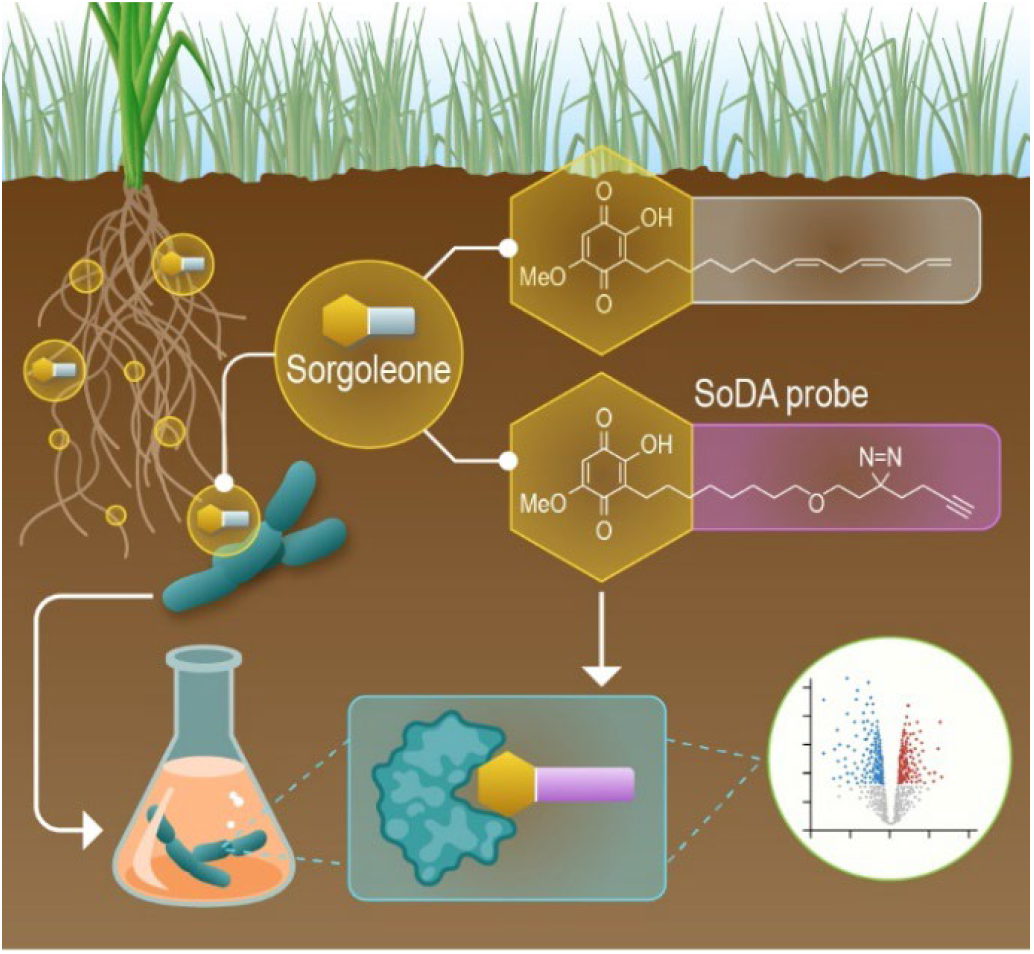
Overview of SoDA-PAL workflow. Sorgoleone (indicated by gold circles) is exuded by sorghum where it comes into contact with rhizosphere bacteria (*A. pittii*, teal bacteria). The structure of sorgoleone informed the design of the SoDA-PAL probe (structures framed by white and pink boxes respectively). Lysates from cultured *A. pittii* can then be probed via SoDA-PAL, leading to the identification of direct sorgoleone protein interactors.

## Results

### The design and synthesis of SoDA-PAL probe

Sargent and Wangchareontrakul previously reported the total synthesis of sorgoleone as a 22-step convergent synthesis, with 15 steps in the longest linear sequence.^46^ We opted for late-stage installation of the diazirine (photo-crosslinking reactive group) and alkyne (“click” tag for attachment of reporter groups) to accommodate the potentially harsh oxidation conditions described for synthesis of the methoxy benzoquinone group. Synthesis of SoDA-PAL was accomplished over 13 total steps in 6% overall yield (**Scheme 1**). We opted for a milder Zemplen deacetylation reaction rather than the lithium aluminum hydride (LAH)-mediated deacetylation as reported in the original total synthesis of sorgoleone. Notably, the penultimate dihydroquinone compound was sensitive to oxidation by ambient air and could not be isolated in pure form. This intermediate was readily converted to the benzoquinone product 11, which has similarly been observed for sorgoleone, which is biosynthesized and released from sorghum roots as the dihydroquinone.^7^

**Scheme 1.**
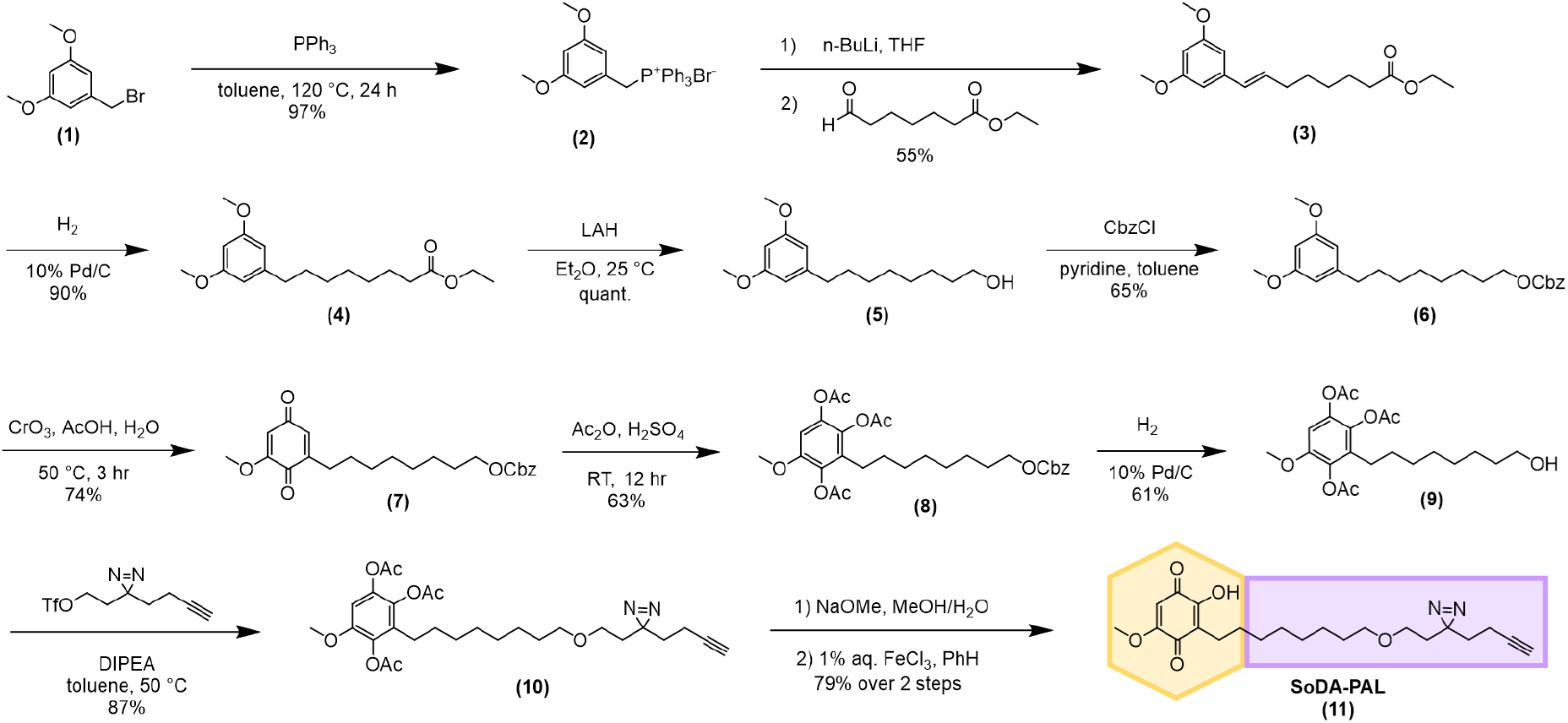
SoDA-PAL probe synthesis.

### Global proteomic analyses reveal carbon source-dependent shifts in the proteome

The sorghum rhizosphere associated bacterium *Acinetobacter pittii* SO1 was found to utilize sorgoleone as a carbon source and was selected as an ideal bacterium for this study.^18^ We evaluated three different culturing strategies that reflect *A. pittii* growth on acetate, sorgoleone, or both (started on acetate then switched to sorgoleone later in the growth stage). Differences in the proteomes of these three culturing conditions (“acetate”, “sorgoleone”, and “acetate to sorgoleone”, respectively) were quantitatively assessed by performing global proteomic analysis of lysates from the three culturing conditions. Global proteomics revealed significant differences in protein abundances between “acetate” cultures versus “sorgoleone” as well as an observable transition in protein abundances in the “acetate to sorgoleone” cultures. Overall, the global proteomes of “acetate to sorgoleone” cultures had the largest number of shared proteins with both “acetate” and “sorgoleone” cultures and the fewest unique proteins (**Figure S3**). These results indicate the occurrence of a carbon-source dependent shift in protein abundances and supported the utilization of the “acetate to sorgoleone” SO1 cultures for downstream SoDA-PAL analysis of *A. pittii* transitioning from growth on different substrates. The global proteomics analysis revealed unique and shared increased peptide coverage related to transport functions across all three growth conditions (A/AS/S). Clustering of transporter proteins reveals protein expression level changes to nutrient transport mechanisms such as ATP-binding cassette (ABC), resistance-nodulation-cell division (RND), multi-drug (MDR), lipopolysaccharide (LPS), and secondary metabolite transporters (**Figure S4**).

### Qualitative evaluation of the SoDA-PAL probe shows dose-dependent and competition-sensitive labelling of Acinetobacter pittii SO1 lysates

As an initial evaluation of the efficacy of the SoDA-PAL probe and to determine optimal probe-protein affinity concentrations, clarified lysates from *A. pittii* SO1, cultured under the “acetate” and “sorgoleone” conditions, were treated with increasing concentrations of the SoDA-PAL probe. Probe-labeled proteins were resolved using sodium dodecyl-sulfate polyacrylamide gel electrophoresis (SDS-PAGE, **Figure 2A**). Increasing concentrations of SoDA-PAL resulted in an increasing fluorescent signal from distinct protein bands (**Figure 2B**), particularly for lower molecular weight proteins. Subtle differences could be observed between labeled proteins from cultures cultured with either acetate or sorgoleone.

**Figure 2.**
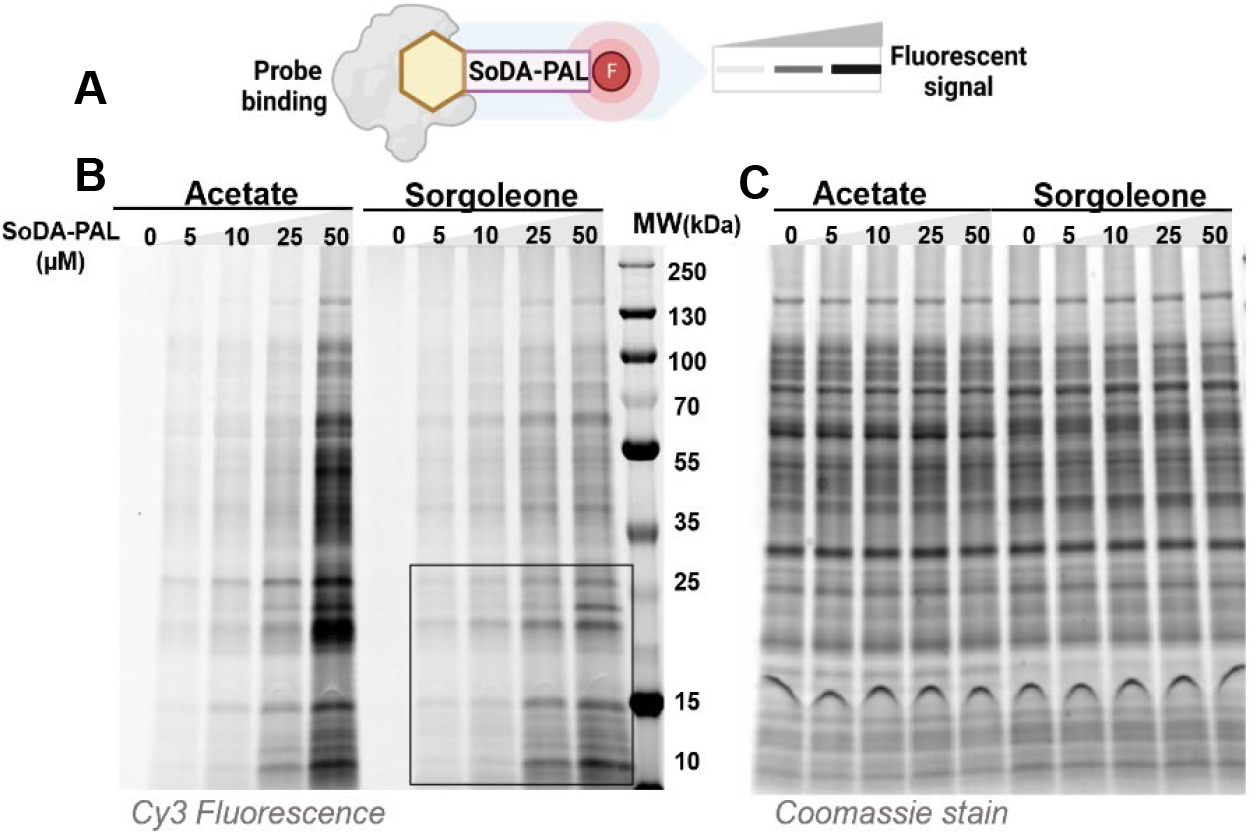
In-gel fluorescence analysis of SO1 clarified lysate cultured under either the “acetate” or “sorgoleone” culturing condition. A. SoDA-PAL probe conjugated with a fluorophore will result in fluorescently labeled target protein. B. SO1 clarified lysates treated with either DMSO (-) or an increasing amount of SoDA-PAL probe (5-50 µM). C. Coomassie stain of probe-labeled gel to indicate equal protein loading.

Importantly, the proteins that were most intensely labeled with the probe were not also the most abundant proteins in the sample, as determined through comparison with the total protein Coomassie stain (**Figure 2C**). An ideal concentration of probe (25 µM) was determined from this experiment and used in later proteomic enrichment experiments.

Additionally, to qualitatively determine the selectivity of the probe for sorgoleone-binding proteins, we performed competition assays by incubating SO1 lysates with increasing concentrations of sorgoleone. Pretreatment of proteins with an excess of sorgoleone results in sorgoleone-specific binding sites being occupied; the SoDA-PAL probe is therefore outcompeted by the sorgoleone, and fewer proteins are labeled. This results in a lower fluorescent signal from the probe when the lysates are analyzed with SDS-PAGE **(Figure 3A)**. The “acetate to sorgoleone” culturing condition was selected for this initial competition evaluation as it exhibits a proteomic profile similar to the “sorgoleone” growth condition

**Figure 3.**
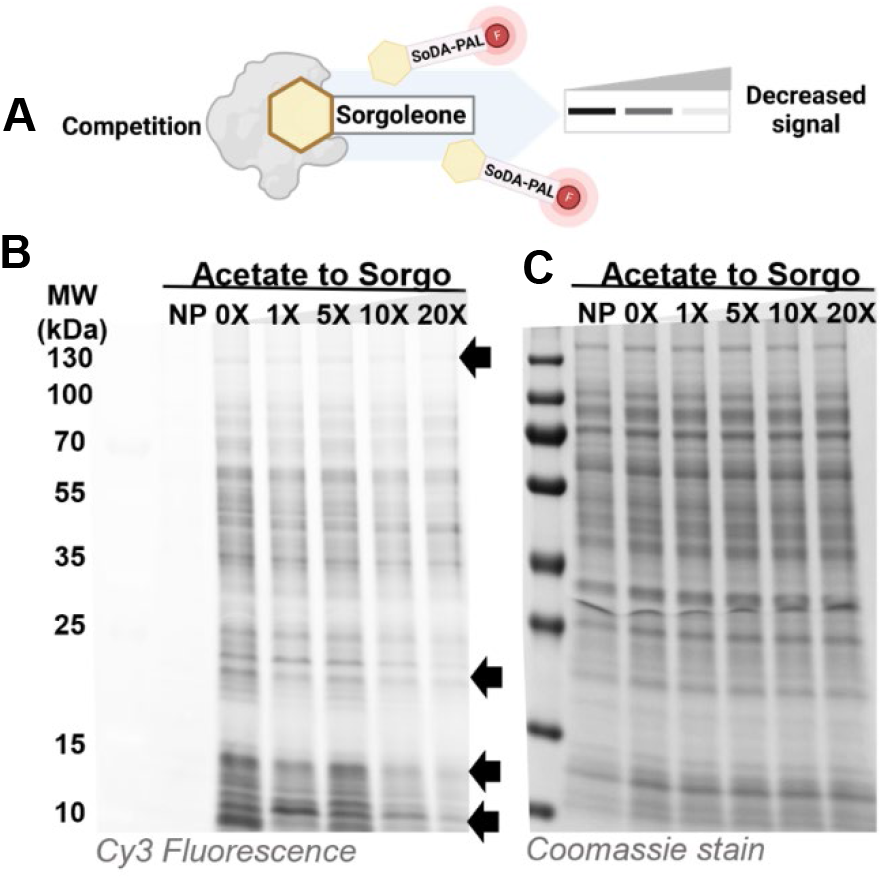
Competition analysis of SO1 clarified lysate pre-treated with sorgoleone. A. For competition analysis, lysates are pre-treated with the competitor (sorgoleone) followed by incubation with SoDA-PAL probe. The sorgoleone outcompetes the SoDA-PAL probe for protein binding sites leading to a decrease of fluorescent signal. B. In-gel fluorescence analysis of SO1 clarified lysate with either DMSO (0X) or a range of competitor (0X, 1X, 5X, 10X, 20X corresponding to 0, 25, 125, 250, 500 µM sorgoleone respectively) followed by probe labelling. Arrows indicate loss of fluorescent signals in competed lanes. C. Coomassie stain of probe-labeled gel to indicate equal protein loading.

**(Figure S1)** yet yielded a much higher biomass of *A pittii*, making this condition preferable for competitive PAL and enrichment. In this evaluation, we observed that probe labelling of specific proteins could be blocked by competition with sorgoleone (**Figure 3B, C**) and again, competition was most readily observed in lower molecular weight proteins.

### SoDA-PAL enriches a subset of the proteome and enables identification of sorgoleone-interacting proteins

After qualitatively confirming successful labeling and competition with sorgoleone in SO1 lysate by SDS-PAGE, we employed a competitive PAL and enrichment protocol followed by mass spectrometry analysis to identify protein targets in the “acetate to sorgoleone” SO1 lysate. For comparing relative abundance values for identification of unique peptide/protein identifications across growth treatment groups, including statistical analysis against no probe (NP) background controls, both proteinGroups.txt and peptides.txt MaxQuant output files were further evaluated for quantitative and qualitative assessment of identified SoDA-PAL protein targets.

We identified 866 proteins as statistically significant for samples labeled with the SoDA-PAL probe compared to the “no probe” (NP) negative control. Protein functions represented by these 866 protein targets included a large number of transporters as well as proteins involved in secondary metabolism and lipid metabolism (**Figures S5, S7**). Sorgoleone competition reduced labeling by SoDA-PAL in 137 of the proteins to a statistically significant extent, representing protein targets that we anticipate are most likely direct binders of sorgoleone; 11 of these proteins were not observed in any global proteomics dataset.

Among these 137 proteins, we identified 19 putative transcriptional regulators, 10 putative transporters, and 8 putative hydrolases; 22 proteins were hypothetical proteins for which no predicted functional annotations were available **(Figure 4, Table S2)**.

**Figure 4.**
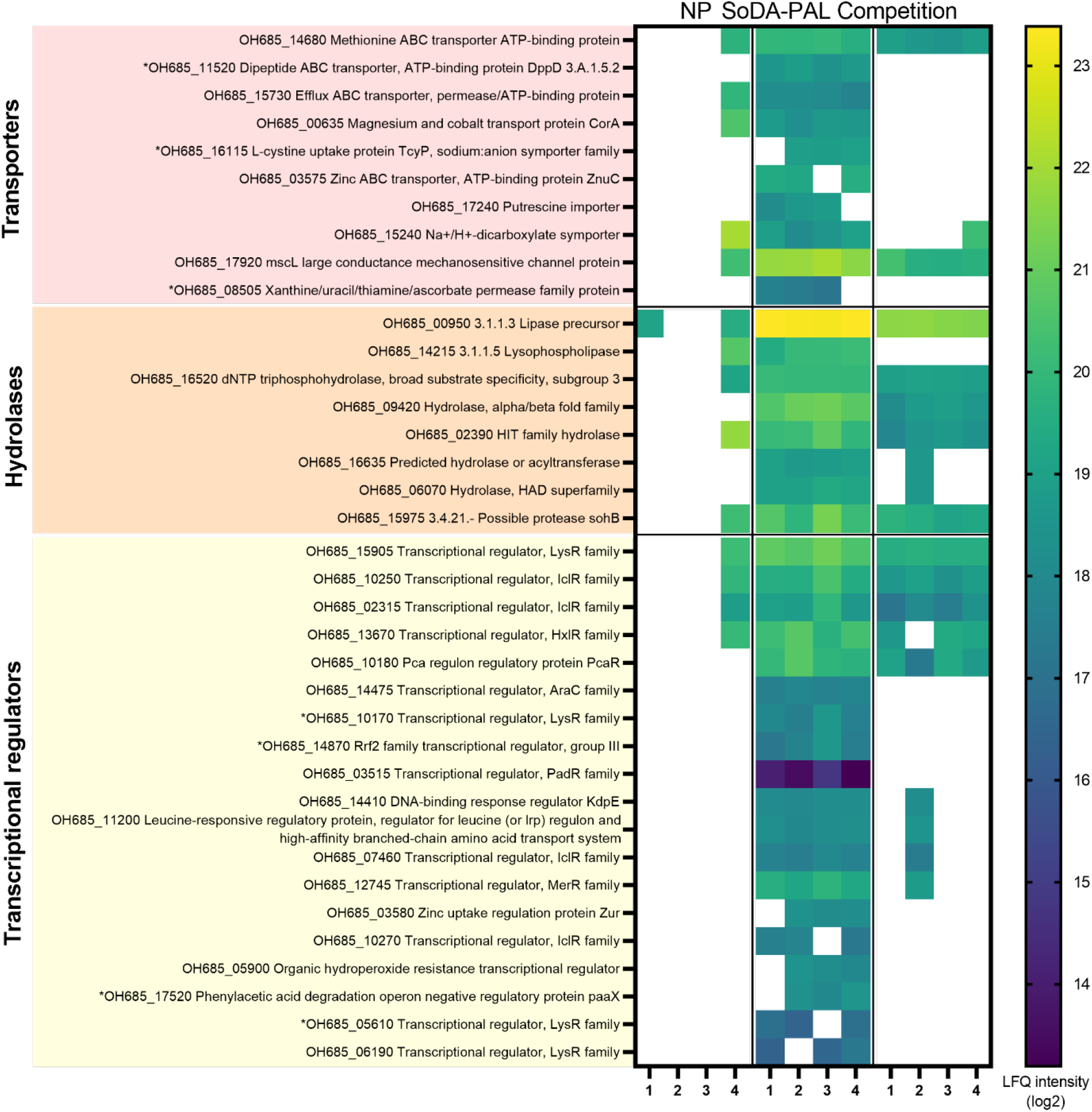
Heatmap of selected sorgoleone protein targets identified in *A. pittii* cultured on acetate to sorgoleone using SoDA-PAL. Label free quantification (LFQ) intensities for each replicate (n = 4) in the no probe (NP), SoDA-PAL only, and Competition sample groups are shown for protein targets with putative transporter, hydrolase, and transcriptional regulator functions; blank boxes are missing values. Proteins that were unique to profiling with SoDA-PAL and not observed in any global proteomics dataset are denoted with an asterisk.

Observed in the set of significantly competed sorgoleone targets were members of a putative sorgoleone catabolism pathway that was recently identified via differential transcriptomic analysis of *Acinetobacter pittii* strain SO1. This pathway is encoded by a gene cluster containing a putative monooxygenase, two putative α/β fold hydrolases, and a cytosine deaminase (**Figure 5A**).^18^ Initially our global proteomic analysis revealed a marked difference in coverage of this pathway between carbon sources, where the lowest pathway member coverage was observed in the acetate culture (A), with higher coverage observed with the acetate to sorgoleone (AS) culture and sorgoleone only culture (S) **(Figure 5B).** This trend was further reinforced with competitive SoDA-PAL labelling of the acetate to sorgoleone cultures, as competition was observed for all four pathway members with the two putative α/β fold hydrolases (OH685_09415, OH685_09420) exhibiting significant competition, implying a direct interaction of those protein targets with sorgoleone **(Figure 5C).**

**Figure 5.**
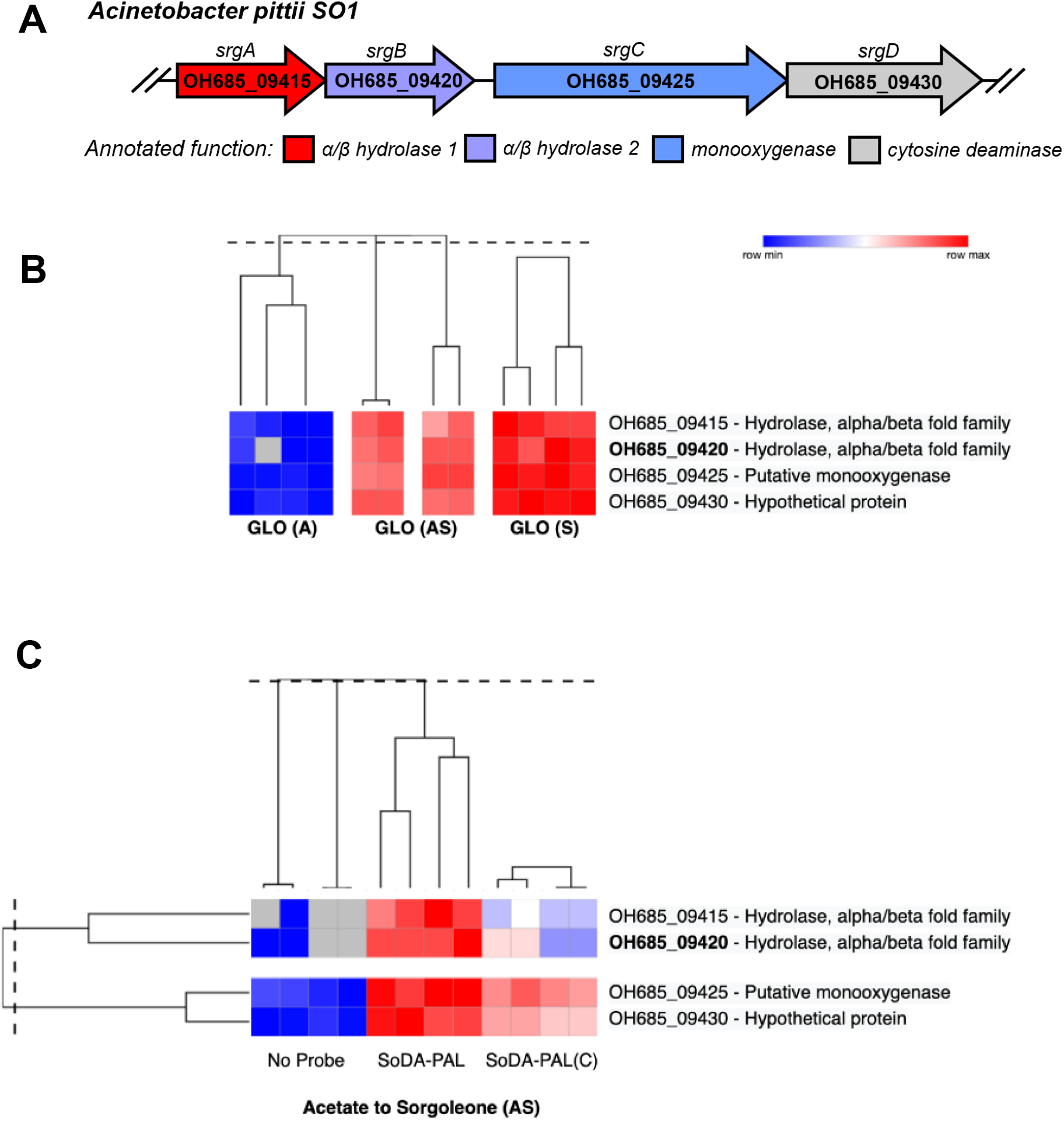
A. Srg cluster identified as putative sorgoleone degradation pathway. B. Global proteomic peptide coverage of Srg gene cluster with increasing protein sequence coverage across the various growth treatments for acetate culture only (A), acetate to sorgoleone (AS) culture, and sorgoleone only culture (S). B. Acetate to sorgoleone growth only SoDA-PAL labeling with sorgoleone competition (C) demonstrating a greater reduction of hydrolase affinity protein targets related to our gene cluster of interest.

### SoDA-PAL target OH685_09420 hydrolase potentially possesses a sorgoleone binding pocket

With our competition data supporting the model that putative sorgoleone degradation pathway member OH685_09420 directly interacts with sorgoleone, we employed molecular modeling analysis on protein target OH685_09420 with the intention of finding potential sorgoleone binding sites. For this, a protein structure was generated using the sequence-to-structure machine learning (ML) model, AlphaFold2 **(Figure 6A).**

**Figure 6.**
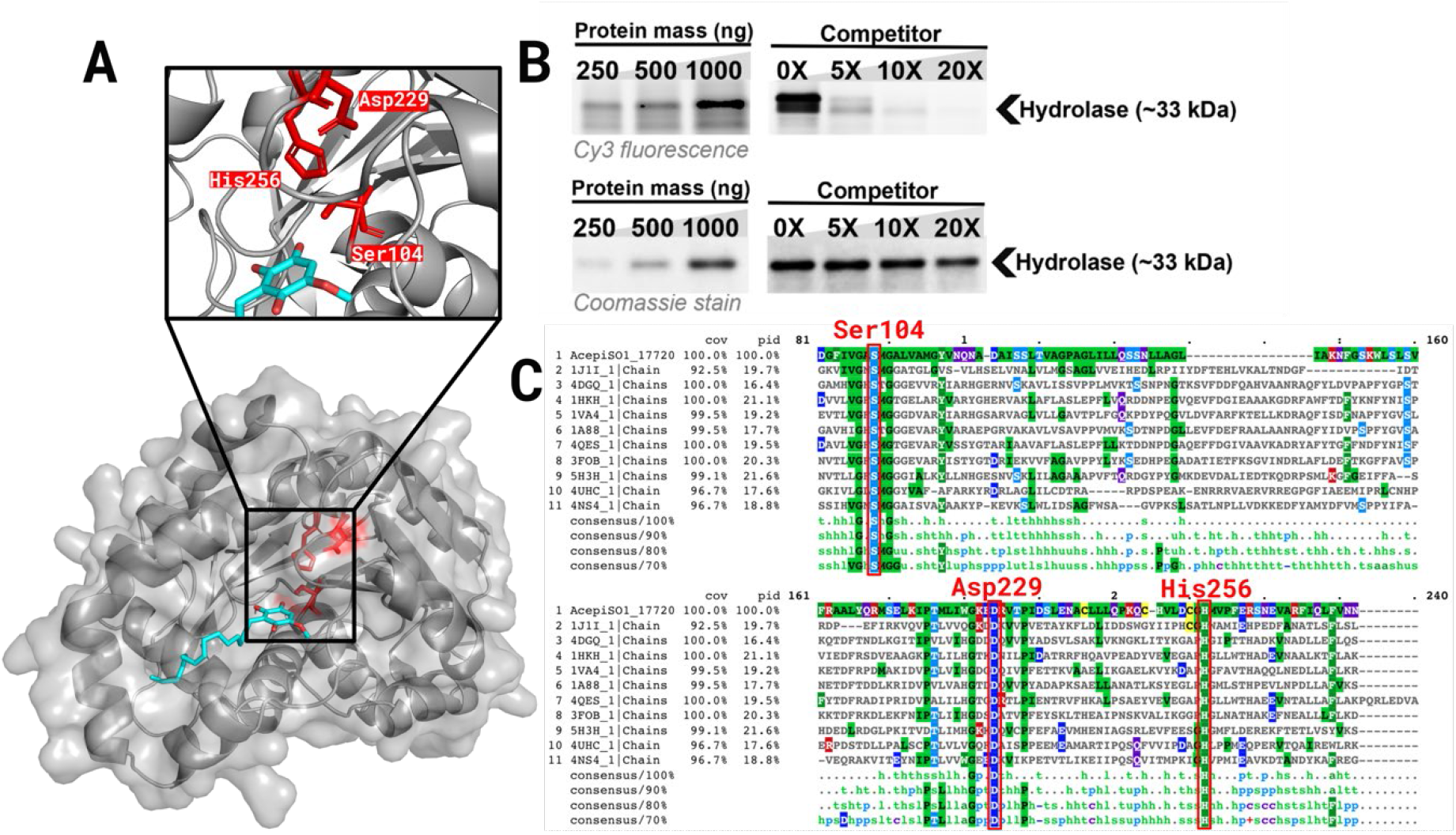
Molecular modeling and in vitro analysis of enriched protein target *OH685_09420*. A. AlphaFold generated model of OH685_09420 (gray structure) with docked sorgoleone model (teal structure) flanked by putative active site residues (red structures). B. In vitro protein labeling (right) and competition assay (left) of recombinantly-expressed OH685_09420. Top panels depict Cy3 fluorescence image of gel, bottom panels depict Coomassie stained gel to show equal protein loading. C. Multiple Sequence Alignment with the top 10 most similar structures to OH685_09420 found through DALI. Conserved residues in putative catalytic triad are boxed in red.

Supporting the genetic pathway annotation, OH685_09420 was classified as a serine hydrolase through structural comparisons to experimentally determined crystal structures found in the Protein Data Bank using DALI. A putative binding site centered on the triad of Ser104, Asp229, and His256 was determined using the ML model p2rank along with a Multiple Sequence Alignment with the top 10 most similar structures found through DALI. This Ser-His-Asp triad is common among serine hydrolases is characterized by the highly nucleophilic serine residue (**Figure 6C**). Identifying the putative pocket and key residues provided us with the information required to run targeted docking studies on the predicted binding site. Two sets of docking studies were performed, one using the SoDA-PAL probe and another with sorgoleone. Each set generated 25 poses within a 20Å cube centered on the catalytic serine. From this, we found the top scoring poses oriented with the benzoquinone group pointed towards the catalytic triad and the tail along the open binding pocket on the outside of the protein (**Figure 6A**).

To continue to explore the potential role of OH685_09420 as a direct sorgoleone binder, we evaluated SoDA-PAL probe binding and direct competition of the probe with sorgoleone in recombinantly expressed and purified OH685_09420. With this recombinantly expressed protein, we observed probe labelling down to 250 ng protein. To determine the binding specificity of the probe for a putative sorgoleone binding site of OH685_09420, we performed a competition assay wherein the recombinant OH685_09420 was incubated in the presence of increasing concentrations of sorgoleone, followed by probe labelling. Competition of SoDA-PAL was immediately evident at a 5X concentration of sorgoleone to probe, and at 20X concentration of sorgoleone, probe labeling was almost entirely competed (**Figure 6B**, Figure S2). This finding reinforces that SoDA-PAL labelling can directly identify sorgoleone binders.

## Discussion

The release of specific metabolites from plant roots, or root exudation, drives plant-microbe interactions, but the molecular pathways by which root exudates shape the sorghum rhizosphere microbiome require further elucidation. To address this, we aimed to design and synthesize a novel photoaffinity probe based on the structure of sorgoleone. The probe could then be used to identify proteins that interact with this sorghum-produced secondary metabolite, with potential future extension of this chemical tool toward profiling complex soil microbial communities for sorgoleone-catabolizing species.

To generate a sorgoleone probe for identifying proteins involved in sorgoleone recognition, transport, and catabolism, we envisioned incorporation of the diazirine photo-crosslinking reactive group^34, 35^ and an alkyne for flexible downstream click chemistry functionalization, into the overall structure of sorgoleone. Notably, the Dayan group has previously characterized a variety of naturally-produced sorgoleone analogs featuring modest diversity in the lipid-like chain.^47^ We therefore hypothesized that modification of this region of sorgoleone would generate a probe molecule that was more likely to be tolerated as a sorgoleone analog compared to probe structures that would involve modification of the benzoquinone group. Treatment of SO1 lysate with SoDA-PAL revealed competition-sensitive labelling, indicating that use of SoDA-PAL for enrichment and proteomic analysis could likely be used to identify sorgoleone-specific protein targets.

Global proteomics results showed that all four gene members of the Srg gene cluster implicated in sorgoleone catabolism were readily identified and were more abundant in culturing conditions with sorgoleone compared to acetate.^18^ From this pathway, OH685_09420 was identified as a protein binder of sorgoleone through competitive labeling with SoDA-PAL and purified sorgoleone. Recent technological advances in structural bioinformatics software, which has enabled faster and more accurate protein structure prediction through artificial intelligence, provide new opportunities to combine experimental and computational approaches for elucidation of protein functions and interactions.^48–50^ To this end, we generated a 3D protein structure for OH685_09420 for protein-ligand docking studies and from this, a putative binding site could be determined along with probe-protein interactions using our computational workflow. The presence of a sorgoleone binding site in this protein was then confirmed by in vitro competition analysis of the recombinant protein target. These experimental and computational results support direct interaction of OH685_09420 with sorgoleone as a potential substrate, whereas the other proteins in the Srg cluster may play more important roles in downstream interactions with intermediate species during sorgoleone catabolism. Thus, SoDA-PAL provides unique insight into the specific molecular interactions between sorgoleone and a subset of the *A. pittii* proteome that complements transcriptomic and global proteomics approaches. Furthermore, proteins were observed in SoDA-PAL-enriched samples that were not identified through global proteomics analyses, highlighting the ability of our ABPP approach to selectively enrich low abundance proteins yet with high affinity for sorgoleone. These types of protein-ligand interactions may not be readily reflected in measurements that depend primarily upon protein or transcript abundance, such as global proteomics and transcriptomics analyses. We also observed a significant enrichment of transcriptional regulators and transporters by SoDA-PAL, including several proteins related to multi-drug resistance and efflux functions (Figures S5-S7). This may be attributed to intrinsic toxicity of quinones, which can act as both electrophiles and electron carriers. The antimicrobial potential of various benzoquinones has been evaluated by multiple research groups.^51, 52^ The influence of quinones on bacterial physiology and metabolism has been reviewed in detail by Franza and Gaudu, including their roles in redox activity, gene regulation, and stress resistance.^53^ Consistent with these previous observations of bacterial responses toward quinones, the SoDA-PAL probe identified two kinases, including one annotated as a putative sensory histidine kinase (OH685_18170) that may respond to environmental cues.^54^ A large number of putative transcriptional regulators were also identified. The presence of sorgoleone, a lipid benzoquinone, may therefore trigger various cellular signaling pathways to mitigate toxicity.^55, 56^ Quinone-sensing transcriptional regulators have been identified in various bacteria,^57^ and RNA-Seq data for SO1 cultured in the presence of sorgoleone was consistent with upregulation of genes associated with stress responses.^18^ Our initial results from the SoDA-PAL probe may provide the basis for opportunities to experimentally confirm binding interactions between sorgoleone and selective transcriptional regulators responsible for sorgoleone-mediated impacts on protein expression which could be used to engineer PGPB for sorghum in the future.

Chemoproteomic approaches such as PAL provide the essential first step for discovery of protein-metabolite interactions by using chemical probes that directly target protein activity or binding to reveal pathways and interactions in complex systems that are invisible to transcriptomic analysis alone. When informed by RNA-Seq data and protein structural modeling, we can further hone PAL proteomic datasets down to a specific active site, enabling efficient prioritization of highly significant proteins for further experimental studies. We have demonstrated this comprehensive workflow, from protein discovery to validation, for the sorghum secondary metabolite sorgoleone in *A. pittii* SO1. Future efforts will further explore these protein targets identified by the SoDA-PAL probe, as well as expand application of the probe to increasingly complex samples, such as mixed microbial isolates. We envision that this workflow will expand our understanding of the sorghum root exudate interactome and elucidate the molecular mechanisms by which specific metabolite shape the sorghum rhizosphere microbiome.

## Data Availability

Primary liquid chromatography mass spectrometry (LC-MS) raw measurement data reported in this study are accessible at the Mass Spectrometry Interactive Virtual Environment (MassIVE) community repository (https://massive.ucsd.edu/), and are available for download under the accession: MSV000091986 (https://massive.ucsd.edu/ProteoSAFe/dataset.jsp?accession=MSV000091986). In efforts to enable dataset citation discovery, transparency, and reproducibility, we ask that the following project digital data assets listed at the PNNL DataHub institutional repository and corresponding domain database accessions be included. Raw measurement proteomic data can be referenced under the digital object identifier (DOI): <u>10.25345/C5P55DS70</u>. All other supporting material, experimental metadata, and processed MaxQuant analysis results reported here can accessed under the dataset drop-down options from the Persistence Control of Engineered Functions in Complex Soil Microbiomes (PerCon SFA), Secure Biosystems Design Project Data Catalog at PNNL DataHub page (https://doi.org/10.25584/1969551), under “<u>DATASET-Proteomics SoDA-PAL Protein Profiling Proteomics of Sorgoleone Catabolizing Microbial Isolate Acinetobacter pittii SO1; LC-MS Proteomics</u>.”

## Funding Acknowledgements

The provided material is based upon work supported by the U.S. Department of Energy, Office of Biological and Environmental Research, Genomic Science Program (https://genomicscience.energy.gov/) and is a contribution of the Pacific Northwest National Laboratory Persistence Control of Engineered Functions in Complex Soil Microbiomes project. A portion of this research is supported under the EMSL user project award <u>10.46936/cpcy.proj.2022.60649/60008751</u>, for leveraging instrument capabilities operating at the Environmental Molecular Sciences Laboratory, a DOE Office of Science User Facility, sponsored by the Biological and Environmental Research program under Contract No. **DE-AC05-76RL01830**.

## Materials and Methods

### SoDA-PAL Probe Synthesis

The photoaffinity sorgoleone diazirine alkyne (SoDA) probe **(11)** was synthesized over 13 steps (Scheme 1). The bromomethyl dimethoxy starting material **(1)** was reacted with triphenylphosphine to afford the phosphonium bromide salt **(2)**. The phosphonium bromide salt was treated with n-BuLi to form the corresponding ylide for a subsequent Wittig reaction with the aldehyde. The resulting alkene **(3)** was hydrogenated over 10% Pd/C to give the ester **(4)**. Reduction of the ester **(4)** using LAH gave the alcohol **(5)**, which was protected by its benzyloxycarbonyl derivative **(6)** and oxidized to yield the quinone **(7)**.

Acetoxylation of the quinone **(7)** gave the triacetoxy product **(8)**. Deprotection of this compound by hydrogenolysis gave the alcohol **(9)**. Triflation of diazirine linker gave alkyne, which was used to alkylate the alcohol to furnish compound **(10)**. Treatment of the alkylated triacetoxy compound **(10)** with sodium methoxide, followed by treatment with aqueous FeCl_3_ provided the final probe **(11)**. Detailed methods and characterization data for synthetic intermediates and products are provided in the supplemental material.

### Sorgoleone Purification

Seeds from the sorghum x sudangrass hybrid SX-19 (Helena Agri-Enterprises) were soaked in sterile deionized water for 4 hr. and then spread onto plastic mesh sheets (85 g dry seeds per sheet) suspended over water in shallow plastic trays.^4^ The seeds were covered with damp paper towels. Trays were covered with aluminum foil to limit light exposure and maintained at 20 °C in a reach-in plant growth chamber or on the benchtop at room temperature. Paper towels were re-wetted with sterile water every 24-48 hr. After 5-10 days of growth, seedling roots were excised, collected in a beaker, and then washed with HPLC grade chloroform for 2 min to extract hydrophobic compounds. The resulting root exudate chloroform solution was filtered and then dried under reduced pressure to a brown residue. Using this method, approximately 14-16 mg crude root exudate per gram root dry mass was obtained (n=18).

The crude root exudate material was purified using a Biotage Isolera purification system for automated flash silica gel column chromatography. Crude root exudates were dissolved in chloroform or dichloromethane and loaded onto pre-packed silica gel columns. Pure sorgoleone was eluted using a gradient of 1-5% MeOH/dichloromethane. Product fractions were combined and dried under reduced pressure to afford pure sorgoleone as a bright orange solid. Sorgoleone identity was confirmed using nuclear magnetic resonance (NMR) spectroscopy and mass spectrometry.^3^ Purified sorgoleone yield was 4.3-9.1 mg per gram of dry root mass (n=18). Purified sorgoleone was dissolved in 100% DMSO for use in microbial cultures and competitive ABPP experiments.

### Bacterial strains and general culture conditions

Frozen stocks (-80 °C) of *Acinetobacter pittii* strain SO1 (GenBank Accession: <u>CP125227.1</u>) were used to inoculate a 5 mL culture of Luria-Bertani (LB) broth. After culturing overnight at 30 °C, the LB cultures were used to inoculate three 25 mL cultures of Mops-based mineral medium (MME, 9.1 mM K2HPO4, 20 mM Mops, 4.3 mM NaCl, 20mM NH_4_Cl, 0.41 mM MgSO_4_, 68 μM CaCl_2_, and 1× MME trace minerals (pH adjusted to 7.0 with KOH). MME trace mineral stock solution (1000×) contains per liter 1 ml of concentrated HCl, 0.5 g of Na_4_EDTA, 2 g of FeCl_3_, and 0.05 g each of H_3_BO_3_, ZnCl_2_, CuCl_2_⋅2H_2_O, MnCl_2_⋅4H2O, (NH_4_)2MoO_4_, CoCl_2_⋅6H_2_O, and NiCl_2_⋅6H_2_O and 20 mM acetate at a 1/100 dilution. These incubated for an additional 36 hours at 30 °C until the cultures reached saturation (OD of 1-1.2). At this point, these cultures were used to inoculate three separate cultures with unique carbon sources for probe analysis. Growth medium conditions consisted of 25 mL MME supplemented with either i) 3 mM sorgoleone only (prepared from a 200 mM stock in DMSO; sorgoleone, “S”), ii) 20 mM acetate only (acetate, “A), or iii) 20 mM acetate transitioned to 1 mM sorgoleone (acetate to sorgoleone, “AS”). Cultures were incubated at 30 °C for 24 hours, after which the “acetate to sorgoleone” culture was pelleted, washed once with MME, and resuspended in 500 mL of MME with 1 mM sorgoleone. The cultures incubated for an additional 48 hours at 30 °C. After incubation, the OD of all the cultures ranged from 1-1.2. The cultures were pelleted by centrifugation and resuspended in Tris-buffered saline with Tween (TBST, 50 mM Tris, pH7.5, 150 mM NaCl, 0.01% Tween20) and filtered with a 20 µm filter to remove excess sorgoleone. The filtered resuspension was pelleted again, washed once with phosphate-buffered saline (PBS, 137 mM NaCl, 2.7 mM KCl, 8 mM Na_2_HPO4, and 2 mM KH_2_PO4) and pelleted cultures were stored at −80 °C.

### Photoaffinity Probe-Enabled Fluorescent SDS-PAGE Analysis

Frozen pellet from a 50 mL culture of *Acinetobacter pittii* strain SO1 were resuspended in 1 mL of PBS and lysed via sonication (37% amplitude, 30 sec cycle, 3 cycles with 20 sec on ice in between cycles). The lysate was clarified via centrifugation (12,000 × g, 10 min at 4 °C) transferred to a fresh microcentrifuge tube for labelling. Protein concentration was determined by bicinchoninic acid (BCA) assay and lysate was normalized to 1 mg/mL for competition assays and labelling.

Increasing concentrations of sorgoleone solubilized in DMSO were added to normalized cell lysate (40 µL per sample) at 0, 25, 125, 250, 500 µM final concentration sorgoleone. These solutions were incubated for 30 min at 25 °C with shaking at 500 *× g*. After incubation was complete, SoDA-PAL (25 µM final concentration) was added to appropriate samples. The samples were covered in foil and incubation continued for another 30 min under the previous conditions. After incubation, samples were transferred to a UV-compatible 96-well plate, and the samples were irradiated for 7 min on ice with a 254 nm wavelength UV-A lamp.

For SDS-PAGE analysis, copper-catalyzed click chemistry was used to attach a fluorophore to probe-labeled proteins for visualizing proteins by fluorescence imaging. To each sample, TAMRA azide (in DMSO; final concentration 30 µM) followed by 2.5 mM sodium ascorbate (freshly prepared), 1 mM tris(3-hydroxypropyltriazolylmethyl) amine (THPTA), and 2 mM CuSO_4_ were sequentially added. The samples were incubated 1 h at 37 °C with shaking at 500 *× g* covered with foil. Afterwards, 2X Tris-Gly sample buffer with 10X NuPAGE reducing agent were added to the samples, and they were subsequently heated at 85 °C for 2 minutes. Samples (20 µL per well) were resolved on a 1.0 mm SDS-PAGE 10-20% Tris-Gly gel at 125 V for 1.5 hours and visualized by in-gel fluorescence on a Typhoon FLA 9500 laser scanner.

### Photoaffinity Probe-Enabled Proteomics Sample Preparation and Enrichment

*Acinetobacter pittii* strain SO1 whole cell lysates were prepared as described previously. Lysate sample stocks (700 μL) were normalized to 2 mg/mL protein concentration using PBS. For competed samples, sorgoleone (500 μM from a 50 mM stock solution in DMSO) or DMSO (1% v/v final concentration) was added, and the samples were incubated for 30 min at 30 °C with agitation. SoDA-PAL probe (3.5 μL of a 5 mM stock for 25 μM final concentration) or DMSO vehicle control was added to samples and incubated for 30 min at 30 °C with agitation. After incubation, samples were transferred to a UV-compatible 24-well plate, and the samples were irradiated for 7 min on ice with a 254 nm wavelength UV-A lamp.

Click chemistry was performed using the following reagents for 700 μL of probe-labeled sample: 3 μL of biotin-azide (from 14 mM stock solution in DMSO), 5 μL of sodium ascorbate (from 345 mM stock solution in water), 5 μL of THPTA (140 mM stock solution in water), and 5 μL of CuSO_4_ (275 mM stock solution in water). The samples were then agitated at 37 °C for 1 h.

Precipitation of proteins was accomplished through the addition of 100 mM NaCl to each sample (13 μL of 5 M NaCl in water) followed by 4 volumes of prechilled acetone (2.6 mL). These samples were placed at −70 °C overnight. Samples were centrifuged, the supernatant was discarded, and pellets were air-dried for 30 min. To the samples was added 700 μL of 1.2% SDS in 1× PBS. Samples were heated to 95 °C for 2 min and then probe sonicated for 6 s, 1 s pulses. Samples were centrifuged and transferred to fresh Eppendorf tubes, leaving any residual pellet behind. A BCA assay was used to quantify the protein concentration of each sample. All samples were normalized to 0.76 mg/mL, for a total mass of 533 µg protein per sample. After normalization, streptavidin enrichment and tryptic digestion were performed as previously described.^58^ Peptides (25 μL) were transferred to 200 μL volume AQ brand silanized glass vial inserts in 2 mL glass autosampler vials (MicroSolv) and stored at −20 °C.

### Global Proteomics Sample Preparation and Digestion

A solution of 1 mg/mL clarified lysate was prepared as described above from whole cell pelleted *Acinetobacter pittii* strain SO1 grown in the presence of acetate “A”, acetate to Sorgoleone “AS”, and/or Sorgoleone “S” as the main carbon source. Each 100 μL sample was reduced by first adding powdered urea to a final concentration of 8 M to each sample followed by DTT (5 mM final concentration). The samples were incubated at 60 °C for 30 min. Samples were then transferred to 3 mL cryovials and diluted 10-fold with 100 mM NH4HCO3 for a final volume of 1.4 mL CaCl_2_ (1 mM final concentration) was added to each sample, followed by addition of LC-MS grade trypsin protease (Thermo Scientific, PI90057) at a 1:50 w/w enzyme to-protein ratio. Samples were digested for 3 h at 37 °C. 1 mL Solid Phase Extraction (SPE) C-18 columns (MilliporeSigma™ Supelco™ Discovery™ DSC-18 SPE Tube, Bed Wt. 50 mg) were used to clean digested peptides. The columns were conditioned 3x with 1 mL MeOH (100%). The columns were then equilibrated with 2 × 1 mL aliquots of water with TFA (0.1%). Samples were loaded onto the columns and were subsequently washed 4× with 1 mL H_2_O: ACN (95:5) with TFA (0.1%). Peptides were eluted into DNA lo-bind Eppendorf tubes with ACN:H_2_O (80:20) with TFA (0.1%). Samples were dried using a speedvac concentrator, followed by reconstitution with 25 mM NH_4_HCO_3_ and centrifugation at 10,000 *× g* to remove debris from column. Protein concentration was determined using BCA assay and lysate samples were normalized to 0.1 mg/mL.

### Liquid Chromatography Mass Spectrometry (LC-MS/MS) Data Acquisition

Global and photoaffinity proteomics samples were analyzed using a Waters nanoAcquity ultra performance liquid chromatography (UPLC) system connected to a Q Exactive Plus Orbitrap mass spectrometer (Thermo Scientific, San Jose, CA). Samples were loaded into a precolumn (150 μM i.d., 4 cm length, packed in-lab with Jupiter C18 packing material, 300 Å pore size, 5 μM particle size: Phenomenex, Torrance, CA, USA) using mobile phase A (0.1% formic acid in water). The separation was carried out in a LC column (75 µm i.d., 70-cm column, packed in-lab with Jupiter C18 packing material, 300-Å pore size, 3 µm particle size (Phenomenex, Terrence, USA) at a flow rate of 300 nL/min using a 120 min gradient of 12-45% mobile phase B (acetonitrile + 0.1% formic acid). To prevent carryover, the column was washed with 95% mobile phase B for 5 min and equilibrated with 1% mobile phase B for 20 min before the next sample injection. The mass spectrometer source was set at 2.2 kV and the ion transfer capillary was heated to 250 °C. The data-dependent acquisition mode was employed to automatically trigger the precursor scan and the MS/MS scans. The MS1 spectra were collected at a scan range of 300-1800 m/z, a resolution of 70,000, an automatic gain control (AGC) target of 3E6, and a maximum injection ion injection time of 20 ms. For MS2, top 12 most intense precursors were isolated with a window of 1.5 m/z and fragmented by higher-energy collisional dissociation (HCD) with a normalized collision energy at 30%. The Orbitrap was used to collect MS/MS spectra at a resolution of 17,500, a maximum AGC target of 1e5, and maximum ion injection time of 50 ms. Each parent ion was fragmented once before being dynamically excluded for 30 s, using a parent ion tolerance window of 20 ppm.

### MaxQuant Proteomics Data Analysis

Bacterial photoaffinity-based protein labeled (PAL) and global (GLO) proteomic data analysis methods for identification of Sorgoleone catabolizing protein targets were analyzed independently using MaxQuant (https://www.maxquant.org/) software (MS:1001583; version 2.1.4.0)^59^ for feature detection and subsequent downstream protein/peptide quantification (HMS-HCD-HMSn). Thermo Scientific RAW dataset files (MS:1000563) were grouped by two different sample growth treatments (cultures grown on acetate “**A**” only or cultures grown on acetate then transitioned to Sorgoleone, “**AS**”) and corresponding probe-labeled condition groups (20 μM Sorgoleone photoaffinity-based labeled, “**SoDA-PAL**” lysates; 20 μM Sorgoleone photoaffinity-based labeled lysates in the presence of 500 μM purified Sorgoleone, “**C + SoDA-PAL**”; DMSO vehicle only no-probe, “**NP**” lysate controls. Global sample datasets contained three separate growth condition treatments (acetate “A” growth medium only, acetate transitioned to Sorgoleone “AS” growth medium, and Sorgoleone “S” growth medium only) and were analyzed independently using MaxQuant software in parallel using identical parameter settings. Spectra were searched using a standard LC-MS run type against *Acinetobacter pittii* SO1 (NCBI TaxonID:48296) from project WGS assembly (GCA_029987435.1) predicted sequence annotated collection SO1_protein_2022-10-06.fasta, corresponding to novel GenBank accession: CP125227.1N-Terminal protein acetylation and methionine oxidation were selected as variable modifications for all datasets. Protein identifications were processed using a maximum false discovery rate (FDR) of 0.01 (∼1%). For increased peptide/protein identifications (unique + razor), match between runs (MBR) was applied within an alignment time window of 20 min using a 3 min match window. For analyzed unique razor peptide quantification results, a minimum of at least one peptide fragment was required to be matched to a single unique protein sequence assignment contained in corresponding proteome file (“S01_protein.fasta”) and a minimum peptide length of 7 amino acids observed across 50% of the technical replicates within each group for continued follow-on analyses (MS:1001833). All additional MaxQuant parameters were ran at software default entries. Relative intensity Based Absolute Quantification (iBAQ) values were log_2_ transformed using the “proteinGroups.txt” output file and required unique peptide fragments to match to a single protein sequence in fasta file (protein count = 1). Distinct label-free peptide spectrum MS/MS counts (MS:1001904) were analyzed using the “peptides.txt” output file and required a minimum of at least one unique peptide count matched to a single protein sequence for included summation results. Integrative Omics Protein Digestion Simulator tool (https://pnnl-comp-mass-spec.github.io/Protein-Digestion-Simulator/) was used for evaluating digested peptide coverage using predicted normalized elution time (NET) values for comparison of resulting ionizable peptide-protein residues identified in the GLO vs PAL proteome coverages. In addition, hydrophobicity index scores were calculated for comparing unique peptide-protein percent sequence coverage results in resulting output tables (SI Table 1), along with an updated GenBank locus tag (OH685) naming key and current protein collection (A. pittii_SO1_2023-05-16.fasta).

### Statistical Analysis

Protein targets identified in proteomics datasets were evaluated using either a two-tailed t-test or g-test for presence or absence of values to determine statistical significance. The g-test is particularly useful for assessing PAL datasets and other enriched samples, which often contain missing values.^60^ Statistically significant proteins had a p-value < 0.05 for either t-test or g-test when comparing probe to “no probe” (NP) or probe to competition groups (n = 4). For proteins evaluated using a t-test, we also implemented a fold change cutoff of ≥ 2 for comparing LFQ intensities between groups to identify proteins that were enriched to a statistically significant extent.

### Structural Protein Analysis

The computational modeling followed steps of a multi-stage computational pipeline integrating multiple structural analysis tools. The 3D structure prediction of the 3 proteins of interest was completed using Alphafold2^48^ implemented through ColabFold.^49^ The predicted structures were compared with experimentally determined protein structures in Protein Data Bank^61^ through the DALI webserver^62^, where the top 10 results based on Z-score were used for the comparison. Binding pocket locations for each predicted structure was determined using p2rank^63^ by selecting the most probable pocket. These pockets were also compared to the available co-crystalized structures returned by DALI.

For the protein-ligand docking studies, we utilized QVina2^64^ an expansion of AutoDock Vina.^65^ The protein structures were prepared for docking using Reduce^66^ for adding hydrogens to the structure and Open Babel^67^ to generate the Vina compatible structure. The 3D ligand conformers were obtained from PubChem^68^ or generated using RDKit (Ver. 2022_03_4).^69^ Docking was centered around the predicted catalytic triad with a grid box size of 30×30×30 Å, up to 25 poses were generated per ligand. The resulting poses were explored using PyMOL.^70^ Top selected poses were further analyzed using the Protein Ligand Interaction Profiler^71^ to obtain more detailed interaction information. All visualizations were done in PyMol.

### Generation of expression plasmids

The coding sequence for OH685_09420 was codon optimized for E. coli, synthesized by Twist Biosciences and inserted into a pET expression vector via Gibson Assembly using NEBuilder® HiFi DNA Assembly Master Mix (NEB) following NEB protocols. Assembled plasmids were transformed into NEB 5-alpha F’Iq (NEB) and standard chemically competent Escherichia coli transformation protocols were used to construct plasmid host strains. Transformants were selected on LB (Miller) agar plates containing 50 µg/mL kanamycin for selection and incubated at 37 °C overnight. Individual colonies were selected and cultured in LB (Miller) media with 50 µg/mL carbenicillin overnight. From this culture, plasmid DNA (pEVF72) was purified from E. coli using GeneJet plasmid miniprep kit (Thermo Scientific) and sequences of all plasmids were confirmed using Sanger sequencing performed by Eurofins Genomics.

### Protein production and purification

Approximately 50 ng of pEVF72 was combined with BL21(DE3) *E. coli* cells using chemical transformations and the resultant colonies were used to inoculate starter cultures (5 mL) by scraping a swath of cells from a fresh LB agar plate. Starter cultures were grown overnight in 2XYT (no more than 16 hr.) in the presence of kanamycin and were used to inoculate expression cultures of ZY505 media (100 mL, 1:100 dilution) containing kanamycin. Expression cultures were grown with constant shaking at 250 rpm at 37°C until reaching an OD_600_ (optical density at 600 nm) of 1.5, wherein they were induced with a final concentration of 0.1 mM IPTG. Upon induction, the temperature was decreased to 25 °C and cultures incubated for an additional 18 hr. before cell harvesting by centrifugation at 5400 relative centrifugal force (rcf).

Cell pellets were resuspended in a lysis/wash buffer [50 mM tris, 500 mM NaCl, and 5 mM imidazole (pH 7.5)] and lysed via sonication. Cell debris was pelleted at 15,000 rcf for 25 min at 4°C and clarified cell lysate was recovered. To bind His_6_-tagged protein, clarified cell lysate was incubated with 150 μL of Ni-NTA resin at 4°C for 1 hour with rocking. Resin was collected and extensively washed with 50 resin bed volumes (bv) of lysis buffer. Bound protein was eluted from the TALON resin by incubation with 5 bv of elution buffer (lysis buffer with 200 mM imidazole) concentrated to 40 μM and exchanged into 50 mM phosphate, 200 mM NaCl, 15 mM Imidazole with a 3-kDa MWCO Vivaspin spin-concentration filter (GE Health Sciences). Protein concentration was determined by absorbance at 280 nm and flash-frozen with liquid nitrogen and stored at −80°C until needed.

### Fluorescent Gel Analysis of Recombinant Proteins

OH685_09420 protein was diluted to 4 μM in 50 uL of PBS. To this protein solution, either SoDA-PAL probe was added at a final concentration of 4 μM or an equivalent volume of DMSO and the reactions were incubated at 25 °C for 30 min with UV irradiation at 254 nm occurring immediately following probe labeling on ice for 7 min.

Following irradiation, click chemistry mediated attachment was performed with the addition of TAMRA azide (4.8 μM) to probe-labeled proteins followed by the addition of sodium ascorbate (400 μM), THPTA tris(3-hydroxypropyltriazolylmethyl)amine (160 μM), and copper sulfate (320 μM). The samples were incubated 1 h at 37 °C with shaking at 500 *× g* covered with foil. Afterwards, 2X Tris-Gly sample buffer with 10X NuPAGE reducing agent were added to the samples, and they were subsequently heated at 85°C for 2 minutes. Samples (20 µL per well) were resolved on a 1.0 mm SDS-PAGE 10-20% Tris-Gly gel at 125 V for 1.5 hours and visualized by in-gel fluorescence on a Typhoon FLA 9500 laser scanner.

For competitive inhibition studies, each pure protein was pre-treated with increasing concentrations of sorgoleone (5X to 20X probe concentration) and incubated 25 °C for 30 minutes. Following incubation, SoDA-PAL probe (4 μM) was added to each reaction, and the samples were treated and visualized as described above.

## AUTHOR ACKNOWLEDGEMENTS

We thank Professor Franck Dayan for his advice on sorgoleone isolation from sorghum seedlings and providing reference material for analysis. We are grateful to Dr. Chathuri Kombala for her assistance with global proteomics sample preparation and Jennifer Walker for preliminary SDS-PAGE experiments. We also thank Dr. Aaron Odgen, Lucas Webber, and Lydia Griggs for developing and carrying out sorgoleone isolation.

## REFERENCES

1. Hossain MS, Islam MN, Rahman MM, Mostofa MG, Khan MAR. Sorghum: A prospective crop for climatic vulnerability, food and nutritional security. Journal of Agriculture and Food Research. 2022;8. doi: 10.1016/j.jafr.2022.100300.

2. Regassa TH, Wortmann CS. Sweet sorghum as a bioenergy crop: Literature review. Biomass and Bioenergy. 2014;64:348–55. doi: 10.1016/j.biombioe.2014.03.052.

3. Dayan FE, Rimando AM, Pan Z, Baerson SR, Gimsing AL, Duke SO. Sorgoleone. Phytochemistry. 2010;71(10):1032-9. Epub 20100410. doi: 10.1016/j.phytochem.2010.03.011. PubMed PMID: 20385394.

4. Dayan FE, Howell J, Weidenhamer JD. Dynamic root exudation of sorgoleone and its in planta mechanism of action. J Exp Bot. 2009;60(7):2107–17. Epub 20090408. doi: 10.1093/jxb/erp082. PubMed PMID: 19357432; PMCID: PMC2682501.

5. Gonzalez VM, Kazimir J, Nimbal C, Weston LA, Cheniae GM. Inhibition of a Photosystem II Electron Transfer Reaction by the Natural Product Sorgoleone. Journal of Agricultural and Food Chemistry. 1997;45(4):1415–21. doi: 10.1021/jf960733w.

6. Hejli AM, Koster KL. The allelochemical sorgoleone inhibits root H+-ATPase and water uptake. J Chem Ecol. 2004;30(11):2181–91. doi: 10.1023/b:joec.0000048782.87862.7f. PubMed PMID: 15672664.

7. Einhellig FA, Souza IF. Phytotoxicity of sorgoleone found in grain Sorghum root exudates. J Chem Ecol. 1992;18(1):1–11. doi: 10.1007/BF00997160. PubMed PMID: 24254628.

8. Weston LA, Czarnota MA. Activity and Persistence of Sorgoleone, a Long-Chain Hydroquinone Produced by Sorghum bicolor. Journal of Crop Production. 2008;4(2):363–77. doi: 10.1300/J144v04n02_17.

9. Gimsing AL, Baelum J, Dayan FE, Locke MA, Sejero LH, Jacobsen CS. Mineralization of the allelochemical sorgoleone in soil. Chemosphere. 2009;76(8):1041–7. Epub 20090602. doi: 10.1016/j.chemosphere.2009.04.048. PubMed PMID: 19493559.

10. Zhalnina K, Louie KB, Hao Z, Mansoori N, da Rocha UN, Shi S, Cho H, Karaoz U, Loque D, Bowen BP, Firestone MK, Northen TR, Brodie EL. Dynamic root exudate chemistry and microbial substrate preferences drive patterns in rhizosphere microbial community assembly. Nat Microbiol. 2018;3(4):470-80. Epub 20180319. doi: 10.1038/s41564-018-0129-3. PubMed PMID: 29556109.

11. Pang Z, Chen J, Wang T, Gao C, Li Z, Guo L, Xu J, Cheng Y. Linking Plant Secondary Metabolites and Plant Microbiomes: A Review. Front Plant Sci. 2021;12:621276. Epub 20210302. doi: 10.3389/fpls.2021.621276. PubMed PMID: 33737943; PMCID: PMC7961088.

12. Wang P, Chai YN, Roston R, Dayan FE, Schachtman DP. The Sorghum bicolor Root Exudate Sorgoleone Shapes Bacterial Communities and Delays Network Formation. mSystems. 2021;6(2). Epub 20210316. doi: 10.1128/mSystems.00749-20. PubMed PMID: 33727394; PMCID: PMC8546980.

13. Sarr PS, Nakamura S, Ando Y, Iwasaki S, Subbarao GV. Sorgoleone production enhances mycorrhizal association and reduces soil nitrification in sorghum. Rhizosphere. 2021;17. doi: 10.1016/j.rhisph.2020.100283.

14. Sarr PS, Ando Y, Nakamura S, Deshpande S, Subbarao GV. Sorgoleone release from sorghum roots shapes the composition of nitrifying populations, total bacteria, and archaea and determines the level of nitrification. Biology and Fertility of Soils. 2019;56(2):145–66. doi: 10.1007/s00374-019-01405-3.

15. Rimando AM, Dayan FE, Czarnota MA, Weston LA, Duke SO. A new photosystem II electron transfer inhibitor from Sorghum bicolor. J Nat Prod. 1998;61(7):927–30. doi: 10.1021/np9800708. PubMed PMID: 9677276.

16. Rimando AM, Dayan FE, Streibig JC. PSII inhibitory activity of resorcinolic lipids from Sorghum bicolor. J Nat Prod. 2003;66(1):42–5. doi: 10.1021/np0203842. PubMed PMID: 12542343.

17. Rasmussen JA, Hejl AM, Einhellig FA, Thomas JA. Sorgoleone from root exudate inhibits mitochondrial functions. J Chem Ecol. 1992;18(2):197–207. doi: 10.1007/bf00993753. PubMed PMID: 24254909.

18. Oda Y, Elmore JR, Nelson WC, Wilson A, Farris Y, Shrestha R, Garcia CF, Pettinga D, Ogden AJ, Baldino H, Alexander WG, Deutschbauer A, Hurtado CV, McDermott JE, Guss AM, Coleman-Derr D, McClure R, Harwood CS, Egbert R. Sorgoleone degradation by sorghum-associated bacteria; an opportunity for enforcing plant growth promotion. bioRxiv. 2023:2023.05.26.542311. doi: 10.1101/2023.05.26.542311.

19. Rokhbakhsh-Zamin F, Sachdev D, Kazemi-Pour N, Engineer A, Pardesi KR, Zinjarde SS, Dhakephalkar PK, Chopade BA. Characterization of Plant-Growth-Promoting Traits of Acinetobacter Species Isolated from Rhizosphere of Pennisetum glaucum. Journal of Microbiology and Biotechnology. 2011;21(6):556–66. doi: 10.4014/jmb.1012.12006.

20. Umapathi M, Chandrasekhar CN, Senthil A, Kalaiselvi T, Santhi R, Ravikesavan R. Isolation, characterization and plant growth-promoting effects of sorghum [Sorghum bicolor (L.) moench] root-associated rhizobacteria and their potential role in drought mitigation. Arch Microbiol. 2022;204(6):354. Epub 20220531. doi: 10.1007/s00203-022-02939-1. PubMed PMID: 35641831.

21. Rajagopal BS, Rao VR, Nagendrappa G, Sethunathan N. Metabolism of carbaryl and carbofuran by soil-enrichment and bacterial cultures. Canadian Journal of Microbiology. 1984;30(12):1458–66. doi: 10.1139/m84-233.

22. McClure R, Naylor D, Farris Y, Davison M, Fansler SJ, Hofmockel KS, Jansson JK. Development and Analysis of a Stable, Reduced Complexity Model Soil Microbiome. Front Microbiol. 2020;11:1987. Epub 20200826. doi: 10.3389/fmicb.2020.01987. PubMed PMID: 32983014; PMCID: PMC7479069.

23. Schutz V, Frindte K, Cui J, Zhang P, Hacquard S, Schulze-Lefert P, Knief C, Schulz M, Dormann P. Differential Impact of Plant Secondary Metabolites on the Soil Microbiota. Front Microbiol. 2021;12:666010. Epub 20210528. doi: 10.3389/fmicb.2021.666010. PubMed PMID: 34122379; PMCID: PMC8195599.

24. Zegeye EK, Brislawn CJ, Farris Y, Fansler SJ, Hofmockel KS, Jansson JK, Wright AT, Graham EB, Naylor D, McClure RS, Bernstein HC. Selection, Succession, and Stabilization of Soil Microbial Consortia. mSystems. 2019;4(4). Epub 20190514. doi: 10.1128/mSystems.00055-19. PubMed PMID: 31098394; PMCID: PMC6517688.

25. McFedries A, Schwaid A, Saghatelian A. Methods for the elucidation of protein-small molecule interactions. Chem Biol. 2013;20(5):667–73. doi: 10.1016/j.chembiol.2013.04.008. PubMed PMID: 23706633.

26. Yu W, Baskin JM. Photoaffinity labeling approaches to elucidate lipid-protein interactions. Curr Opin Chem Biol. 2022;69:102173. Epub 20220617. doi: 10.1016/j.cbpa.2022.102173. PubMed PMID: 35724595.

27. Chuang HC, Liu MF, Wu HY, Wu YT, Cheng TR, Fang JM. Photoaffinity labeling of benzophenone-containing salicylanilide compounds to give an insight into the mechanism in disrupting peptidoglycan formation. Bioorg Med Chem. 2022;67:116819. Epub 20220513. doi: 10.1016/j.bmc.2022.116819. PubMed PMID: 35635930.

28. Ishikawa F, Konno S, Uchiyama Y, Kakeya H, Tanabe G. Exploring a chemical scaffold for rapid and selective photoaffinity labelling of non-ribosomal peptide synthetases in living bacterial cells. Philos Trans R Soc Lond B Biol Sci. 2023;378(1871):020220026. Epub 20230111. doi: 10.1098/rstb.2022.0026. PubMed PMID: 36633280; PMCID: PMC9835605.

29. Whidbey C, Wright AT. Activity-Based Protein Profiling-Enabling Multimodal Functional Studies of Microbial Communities. Curr Top Microbiol Immunol. 2019;420:1–21. doi: 10.1007/82_2018_128. PubMed PMID: 30406866; PMCID: PMC6561099.

30. Whidbey C, Sadler NC, Nair RN, Volk RF, DeLeon AJ, Bramer LM, Fansler SJ, Hansen JR, Shukla AK, Jansson JK, Thrall BD, Wright AT. A Probe-Enabled Approach for the Selective Isolation and Characterization of Functionally Active Subpopulations in the Gut Microbiome. J Am Chem Soc. 2019;141(1):42–7. Epub 20181217. doi: 10.1021/jacs.8b09668. PubMed PMID: 30541282; PMCID: PMC6533105.

31. Zegeye EK, Sadler NC, Lomas GX, Attah IK, Jansson JK, Hofmockel KS, Anderton CR, Wright AT. Activity-Based Protein Profiling of Chitin Catabolism. Chembiochem. 2021;22(4):717–23. Epub 20201117. doi: 10.1002/cbic.202000616. PubMed PMID: 33049124.

32. Kaschani F, Verhelst SH, van Swieten PF, Verdoes M, Wong CS, Wang Z, Kaiser M, Overkleeft HS, Bogyo M, van der Hoorn RA. Minitags for small molecules: detecting targets of reactive small molecules in living plant tissues using ’click chemistry’. Plant J. 2009;57(2):373–85. Epub 20081025. doi: 10.1111/j.1365-313X.2008.03683.x. PubMed PMID: 18786180.

33. Lin VS. Interrogating Plant-Microbe Interactions with Chemical Tools: Click Chemistry Reagents for Metabolic Labeling and Activity-Based Probes. Molecules. 2021;26(1). Epub 20210105. doi: 10.3390/molecules26010243. PubMed PMID: 33466477; PMCID: PMC7796436.

34. Hill JR, Robertson AAB. Fishing for Drug Targets: A Focus on Diazirine Photoaffinity Probe Synthesis. J Med Chem. 2018;61(16):6945–63. Epub 20180509. doi: 10.1021/acs.jmedchem.7b01561. PubMed PMID: 29683660.

35. Smith E, Collins I. Photoaffinity labeling in target-and binding-site identification. Future Med Chem. 2015;7(2):159–83. doi: 10.4155/fmc.14.152. PubMed PMID: 25686004; PMCID: PMC4413435.

36. Burton NR, Kim P, Backus KM. Photoaffinity labelling strategies for mapping the small molecule-protein interactome. Org Biomol Chem. 2021;19(36):7792–809. Epub 20210922. doi: 10.1039/d1ob01353j. PubMed PMID: 34549230; PMCID: PMC8489259.

37. Rosnow JJ, Hwang S, Killinger BJ, Kim YM, Moore RJ, Lindemann SR, Maupin-Furlow JA, Wright AT. A Cobalamin Activity-Based Probe Enables Microbial Cell Growth and Finds New Cobalamin-Protein Interactions across Domains. Appl Environ Microbiol. 2018;84(18). Epub 20180831. doi: 10.1128/AEM.00955-18. PubMed PMID: 30006406; PMCID: PMC6121979.

38. Balaratnam S, Rhodes C, Bume DD, Connelly C, Lai CC, Kelley JA, Yazdani K, Homan PJ, Incarnato D, Numata T, Schneekloth JS, Jr. A chemical probe based on the PreQ(1) metabolite enables transcriptome-wide mapping of binding sites. Nat Commun. 2021;12(1):5856. Epub 20211006. doi: 10.1038/s41467-021-25973-x. PubMed PMID: 34615874; PMCID: PMC8494917.

39. Chen X, Wang Y, Ma N, Tian J, Shao Y, Zhu B, Wong YK, Liang Z, Zou C, Wang J. Target identification of natural medicine with chemical proteomics approach: probe synthesis, target fishing and protein identification. Signal Transduct Target Ther. 2020;5(1):72. Epub 20200521. doi: 10.1038/s41392-020-0186-y. PubMed PMID: 32435053; PMCID: PMC7239890.

40. Luzarowski M, Skirycz A. Emerging strategies for the identification of protein-metabolite interactions. J Exp Bot. 2019;70(18):4605–18. doi: 10.1093/jxb/erz228. PubMed PMID: 31087097; PMCID: PMC6760282.

41. Li W, Zhou Y, You W, Yang M, Ma Y, Wang M, Wang Y, Yuan S, Xiao Y. Development of Photoaffinity Probe for the Discovery of Steviol Glycosides Biosynthesis Pathway in Stevia rebuadiana and Rapid Substrate Screening. ACS Chem Biol. 2018;13(8):1944–9. Epub 20180612. doi: 10.1021/acschembio.8b00285. PubMed PMID: 29863335.

42. Brinegar AC, Cooper G, Stevens A, Hauer CR, Shabanowitz J, Hunt DF, Fox JE. Characterization of a benzyladenine binding-site peptide isolated from a wheat cytokinin-binding protein: sequence analysis and identification of a single affinity-labeled histidine residue by mass spectrometry. Proc Natl Acad Sci U S A. 1988;85(16):5927–31. doi: 10.1073/pnas.85.16.5927. PubMed PMID: 3413067; PMCID: PMC281878.

43. Manohar M, Tian M, Moreau M, Park SW, Choi HW, Fei Z, Friso G, Asif M, Manosalva P, von Dahl CC, Shi K, Ma S, Dinesh-Kumar SP, O’Doherty I, Schroeder FC, van Wijk KJ, Klessig DF. Identification of multiple salicylic acid-binding proteins using two high throughput screens. Front Plant Sci. 2014;5:777. Epub 20150112. doi: 10.3389/fpls.2014.00777. PubMed PMID: 25628632; PMCID: PMC4290489.

44. Jones AM, Venis MA. Photoaffinity labeling of indole-3-acetic acid-binding proteins in maize. Proc Natl Acad Sci U S A. 1989;86(16):6153–6. doi: 10.1073/pnas.86.16.6153. PubMed PMID: 16594060; PMCID: PMC297795.

45. Hooley R, Smith SJ, Beale MH, Walker RP. In vivo Photoaffinity Labelling of Gibberellin-Binding Proteins in Avena fatua Aleurone. Functional Plant Biology. 1993;20(5). doi: 10.1071/pp9930573.

46. Sargent MV, Wangchareontrakul S. The synthesis of the first natural host germination stimulant for Striga asiatica(witchweed). Journal of the Chemical Society, Perkin Transactions 1. 1990(5). doi: 10.1039/p19900001429.

47. Kagan IA, Rimando AM, Dayan FE. Chromatographic separation and in vitro activity of sorgoleone congeners from the roots of sorghum bicolor. J Agric Food Chem. 2003;51(26):7589–95. doi: 10.1021/jf034789j. PubMed PMID: 14664512.

48. Jumper J, Evans R, Pritzel A, Green T, Figurnov M, Ronneberger O, Tunyasuvunakool K, Bates R, Zidek A, Potapenko A, Bridgland A, Meyer C, Kohl SAA, Ballard AJ, Cowie A, Romera-Paredes B, Nikolov S, Jain R, Adler J, Back T, Petersen S, Reiman D, Clancy E, Zielinski M, Steinegger M, Pacholska M, Berghammer T, Bodenstein S, Silver D, Vinyals O, Senior AW, Kavukcuoglu K, Kohli P, Hassabis D. Highly accurate protein structure prediction with AlphaFold. Nature. 2021;596(7873):583–9. Epub 20210715. doi: 10.1038/s41586-021-03819-2. PubMed PMID: 34265844; PMCID: PMC8371605.

49. Mirdita M, Schutze K, Moriwaki Y, Heo L, Ovchinnikov S, Steinegger M. ColabFold: making protein folding accessible to all. Nat Methods. 2022;19(6):679–82. Epub 20220530. doi: 10.1038/s41592-022-01488-1. PubMed PMID: 35637307; PMCID: PMC9184281.

50. Baek M, DiMaio F, Anishchenko I, Dauparas J, Ovchinnikov S, Lee GR, Wang J, Cong Q, Kinch LN, Schaeffer RD, Millan C, Park H, Adams C, Glassman CR, DeGiovanni A, Pereira JH, Rodrigues AV, van Dijk AA, Ebrecht AC, Opperman DJ, Sagmeister T, Buhlheller C, Pavkov-Keller T, Rathinaswamy MK, Dalwadi U, Yip CK, Burke JE, Garcia KC, Grishin NV, Adams PD, Read RJ, Baker D. Accurate prediction of protein structures and interactions using a three-track neural network. Science. 2021;373(6557):871–6. Epub 20210715. doi: 10.1126/science.abj8754. PubMed PMID: 34282049; PMCID: PMC7612213.

51. Lana EJ, Carazza F, Takahashi JA. Antibacterial evaluation of 1,4-benzoquinone derivatives. J Agric Food Chem. 2006;54(6):2053–6. doi: 10.1021/jf052407z. PubMed PMID: 16536574.

52. Dahlem Junior MA, Nguema Edzang RW, Catto AL, Raimundo JM. Quinones as an Efficient Molecular Scaffold in the Antibacterial/Antifungal or Antitumoral Arsenal. Int J Mol Sci. 2022;23(22). Epub 20221115. doi: 10.3390/ijms232214108. PubMed PMID: 36430585; PMCID: PMC9697455.

53. Franza T, Gaudu P. Quinones: more than electron shuttles. Res Microbiol. 2022;173(6-7):103953. Epub 20220422. doi: 10.1016/j.resmic.2022.103953. PubMed PMID: 35470045.

54. Koler M, Frank V, Amartely H, Friedler A, Vaknin A. Dynamic Clustering of the Bacterial Sensory Kinase BaeS. PLoS One. 2016;11(3):e0150349. Epub 20160307. doi: 10.1371/journal.pone.0150349. PubMed PMID: 26950881; PMCID: PMC4780735.

55. Davies JS, Currie MJ, Wright JD, Newton-Vesty MC, North RA, Mace PD, Allison JR, Dobson RCJ. Selective Nutrient Transport in Bacteria: Multicomponent Transporter Systems Reign Supreme. Front Mol Biosci. 2021;8:699222. Epub 20210629. doi: 10.3389/fmolb.2021.699222. PubMed PMID: 34268334; PMCID: PMC8276074.

56. Ryan A, Kaplan E, Nebel JC, Polycarpou E, Crescente V, Lowe E, Preston GM, Sim E. Identification of NAD(P)H quinone oxidoreductase activity in azoreductases from P. aeruginosa: azoreductases and NAD(P)H quinone oxidoreductases belong to the same FMN-dependent superfamily of enzymes. PLoS One. 2014;9(6):e98551. Epub 20140610. doi: 10.1371/journal.pone.0098551. PubMed PMID: 24915188; PMCID: PMC4051601.

57. Fritsch VN, Loi VV, Busche T, Sommer A, Tedin K, Nurnberg DJ, Kalinowski J, Bernhardt J, Fulde M, Antelmann H. The MarR-Type Repressor MhqR Confers Quinone and Antimicrobial Resistance in Staphylococcus aureus. Antioxid Redox Signal. 2019;31(16):1235–52. Epub 20190809. doi: 10.1089/ars.2019.7750. PubMed PMID: 31310152; PMCID: PMC6798810.

58. Lin VS, Volk RF, DeLeon AJ, Anderson LN, Purvine SO, Shukla AK, Bernstein HC, Smith JN, Wright AT. Structure Dependent Determination of Organophosphate Targets in Mammalian Tissues Using Activity-Based Protein Profiling. Chem Res Toxicol. 2020;33(2):414–25. Epub 20200110. doi: 10.1021/acs.chemrestox.9b00344. PubMed PMID: 31872761; PMCID: PMC9014469.

59. Cox J, Mann M. MaxQuant enables high peptide identification rates, individualized p.p.b.-range mass accuracies and proteome-wide protein quantification. Nat Biotechnol. 2008;26(12):1367–72. Epub 20081130. doi: 10.1038/nbt.1511. PubMed PMID: 19029910.

60. Webb-Robertson BJ, McCue LA, Waters KM, Matzke MM, Jacobs JM, Metz TO, Varnum SM, Pounds JG. Combined statistical analyses of peptide intensities and peptide occurrences improves identification of significant peptides from MS-based proteomics data. J Proteome Res. 2010;9(11):5748–56. Epub 20101008. doi: 10.1021/pr1005247. PubMed PMID: 20831241; PMCID: PMC2974810.

61. Berman HM, Westbrook J, Feng Z, Gilliland G, Bhat TN, Weissig H, Shindyalov IN, Bourne PE. The Protein Data Bank. Nucleic Acids Res. 2000;28(1):235–42. doi: 10.1093/nar/28.1.235. PubMed PMID: 10592235; PMCID: PMC102472.

62. Holm L. Dali server: structural unification of protein families. Nucleic Acids Res. 2022;50(W1):W210–W5. doi: 10.1093/nar/gkac387. PubMed PMID: 35610055; PMCID: PMC9252788.

63. Krivak R, Hoksza D. P2Rank: machine learning based tool for rapid and accurate prediction of ligand binding sites from protein structure. J Cheminform. 2018;10(1):39. Epub 20180814. doi: 10.1186/s13321-018-0285-8. PubMed PMID: 30109435; PMCID: PMC6091426.

64. Alhossary A, Handoko SD, Mu Y, Kwoh CK. Fast, accurate, and reliable molecular docking with QuickVina 2. Bioinformatics. 2015;31(13):2214–6. Epub 20150224. doi: 10.1093/bioinformatics/btv082. PubMed PMID: 25717194.

65. Trott O, Olson AJ. AutoDock Vina: improving the speed and accuracy of docking with a new scoring function, efficient optimization, and multithreading. J Comput Chem. 2010;31(2):455–61. doi: 10.1002/jcc.21334. PubMed PMID: 19499576; PMCID: PMC3041641.

66. Word JM, Lovell SC, Richardson JS, Richardson DC. Asparagine and glutamine: using hydrogen atom contacts in the choice of side-chain amide orientation. J Mol Biol. 1999;285(4):1735–47. doi: 10.1006/jmbi.1998.2401. PubMed PMID: 9917408.

67. O’Boyle NM, Banck M, James CA, Morley C, Vandermeersch T, Hutchison GR. Open Babel: An open chemical toolbox. J Cheminform. 2011;3:33. Epub 20111007. doi: 10.1186/1758-2946-3-33. PubMed PMID: 21982300; PMCID: PMC3198950.

68. Kim S, Chen J, Cheng T, Gindulyte A, He J, He S, Li Q, Shoemaker BA, Thiessen PA, Yu B, Zaslavsky L, Zhang J, Bolton EE. PubChem 2023 update. Nucleic Acids Res. 2023;51(D1):D1373–D80. doi: 10.1093/nar/gkac956. PubMed PMID: 36305812; PMCID: PMC9825602.

69. Landrum GP, T.; Kelley, B.; Ric; sriniker; gedeck; Vianello, R.; Cosgrove, D.; Schneider, N.; Kawashima, E.; N., Dan; Dalke, A.; Jones, G.; Cole, B.; Swain, M.; Turk, S.; Savelyev, A.; Vaucher, A.; Wójcikowski, M.; Take, I.; Probst, D.; Scalfani, V. F.; Ujihara, K.; Godin, G.; Pahl, A.; Berenger, F.; JLVarjo; jasondbiggs; strets123; JP. RDKit: Open-source cheminformatics. https://www.rdkit.org, Zenodo, 10.5281/zenodo.6798971. rdkit/rdkit: 2022_09_4 (Q3 2022) ed2022.

70. The PyMOL Molecular Graphics System, Version 2.0. Schrödinger, LLC.

71. Adasme MF, Linnemann KL, Bolz SN, Kaiser F, Salentin S, Haupt VJ, Schroeder M. PLIP 2021: expanding the scope of the protein-ligand interaction profiler to DNA and RNA. Nucleic Acids Res. 2021;49(W1):W530–W4. doi: 10.1093/nar/gkab294. PubMed PMID: 33950214; PMCID: PMC8262720.

