## Supplementary Information for "Profiling sorghum-microbe interactions with a specialized photoaffinity probe identifies key sorgoleone binders in *Acinetobacter pittii*"

**Supporting Information**

***Synthetic methods:***

Chemical reagents and solvents were purchased from Sigma-Aldrich, Acros, Alfa Aesar, TCI America, Fisher Scientific, and VWR and were used without further purification. Ethyl 7-oxoheptanoate was purchased from FUJIFILM Wako Chemicals U.S.A. Corporation or synthesized from cycloheptanone via a Baeyer-Villiger oxidation[1, 2] and ester hydrolysis, followed by oxidation with TEMPO/BAIB.[3] Cycloheptanone was purchased from Thermo Scientific. 1-(Bromomethyl)-3 5-dimethoxybenzene and 2-(3-but-3-ynyl-3H-diazirin-3-yl)-ethanol were purchased from Ambeed. Dry THF and $CH_2Cl_2$ were obtained from an LC Tech SP-1 Stand Alone Solvent Purification System. Automated flash silica gel column chromatography was performed using a Biotage Isolera purification system. Analytical thin layer chromatography (TLC) was performed using silica gel 60 F254 plates (0.25 mm) and compounds visualized by shortwave UV irradiation with a handheld UV lamp or staining as indicated.

$^{1}$H NMR spectra were acquired in $CDCl_3$ or DMSO-$d_6$ (Cambridge Isotopes, Tewksbury, MA) at ambient temperature (25 °C) on a Bruker 400 MHz Avance III spectrometer equipped with a 5 mm BBFO SmartProbe. All chemical shifts are reported in the standard notation of parts per million using the peak of the residual proton or carbon signal of $CDCl_3$ ($^{1}$H NMR δ 7.26 and $^{13}$C NMR δ 77.36 ppm) or DMSO-$d_6$ ($^{1}$H NMR δ 2.50 and $^{13}$C NMR δ 39.52 ppm) as an internal reference. Splitting patterns are indicated as follows: s, singlet; d, doublet; t, triplet; m, multiplet; dd, doublet of doublets. LRMS-ESI was performed using a Finnigan LTQ MS (Thermo Electron Corporation).

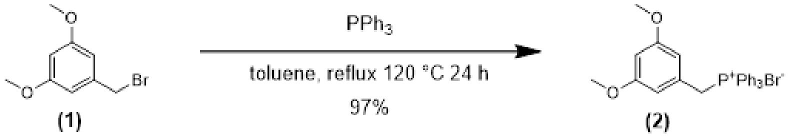


**Scheme S1. Synthesis of (3,5-dimethoxybenzyl)triphenylphosphonium bromide (2)** The phosphonium bromide **(2)** was prepared as previously described.[4] To an oven dried 250 mL flask and stir bar under nitrogen was added 6.00 g (26.0 mmol, 1 eq) of 3,5-dimethoxy benzyl bromide, followed by 60 mL of anhydrous toluene. The mixture was stirred until the solid completely dissolved. 10.2 g (38.9 mmol, 1.5 eq) triphenylphosphine was added to the solution in a single portion. The solution was refluxed for 24 h. After 24 h, the reaction mixture was allowed to cool to room temperature and then the precipitate was removed by filtration. The solids were washed with hexanes, transferred to a glass vial, and dried under vacuum. 12.3 g (96%) of product was collected as a white powder. This material was used in the next step without further purification. MS: m/z = +413 (observed), [M-Br] = 413.5 (calculated). $^{1}$H NMR (400 MHz, DMSO-$d_6$) δ 7.91 (td, $J$ = 7.3, 1.7 Hz, 3H), 7.76 (td, $J$ = 7.8, 3.5 Hz, 6H), 7.72 – 7.64 (m, 6H), 6.42 (q, $J$ = 2.3 Hz, 1H), 6.11 (t, $J$ = 2.4 Hz, 2H), 5.04 (d, $J$ = 15.6 Hz, 2H), 3.50 (s, 6H). $^{13}$C NMR (101 MHz, DMSO -$d_6$) δ 160.38, 160.35, 135.14, 135.10, 134.11, 134.01, 130.16, 130.03, 129.88, 129.79, 118.25, 117.40, 108.92, 108.87, 100.24, 100.20, 55.06, 28.67, 28.20.

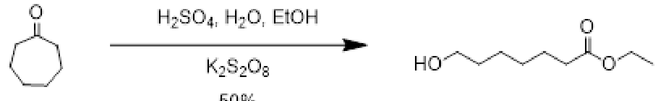


(2a) (2b)

**Scheme S2. Synthesis of ethyl 7-hydroxyheptanoate (2b)** Primary alcohol **(2b)** was synthesized was cycloheptanone via a Baeyer-Villiger oxidation followed by ester hydrolysis, as previously reported.[1, 2] Briefly, conc. sulfuric acid (70 mL), water (25 mL), and absolute ethanol (100 mL) were cooled in a 1 L round bottom flask over wet ice. Potassium persulfate (72 g, 267 mmol 3 eq) was added, portion-wise, with rapid stirring. A solution of cycloheptanone (10.0 g, 89 mmol, 1 eq) in ethanol (30 mL) was added dropwise very slowly to the reaction flask using an addition funnel. The reaction was highly exothermic. The reaction mixture was stirred overnight (20 hr.), warming to r.t. The mixture was diluted with water (300 mL) and extracted with diethyl ether (4 x 100 mL). The combined organic layers were washed 1x with water, dried over $Na_2SO_4$, and concentrated *in vacuo* to a beige oil. The product was purified via flash chromatography on silica gel, eluting 10-40% EtOAc/hex. The product was isolated as a clear oil (7.7 g, 50% yield), $R_f$ (50% EtOAc/hex) = 0.49. $^1$H NMR (400 MHz, $CDCl_3$) δ 4.12 (q, *J* = 7.1 Hz, 2H), 3.64 (t, *J* = 6.6 Hz, 2H), 2.29 (t, *J* = 7.5 Hz, 2H), 1.74 – 1.49 (m, 5H), 1.45 – 1.29 (m, 4H), 1.25 (t, *J* = 7.1 Hz, 3H).

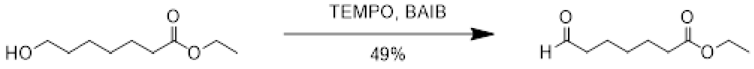


(2b) (2c)

**Scheme S3. Synthesis of ethyl 7-oxoheptanoate (2c).** Aldehyde **(2c)** was synthesized from primary alcohol **(2b)** via a TEMPO-catalyzed oxidation, as previously described.[5] Briefly, ethyl 7-hydroxyheptanoate (2b; 0.53 g, 3.0 mmol, 1 eq) was dissolved in DCM (25 mL), after which (diacetoxyiodo)benzene (BAIB, 1.18 g, 3.65 mmol, 1.2 eq) followed by TEMPO (0.098 g, 0.62 mmol, 0.2 eq) were added. The reaction mixture was stirred at room temperature and monitored by TLC with $KMnO_4$ staining $R_f$ (50% EtOAc/hex) = 0.73. After 16 hr., no starting material was observed by TLC, and the reaction was quenched by addition of sat. $NaHCO_3$ (aq) and sat. $Na_2S_2O_3$ (aq). The product was extracted with DCM, and the combined organic layers were dried over $Na_2SO_4$. The crude material was concentrated *in vacuo* to an orange oil and purified via flash chromatography on silica gel, eluting 5-20% EtOAc/hex. The product was isolated as a clear, very pale orange oil (0.25 g, 49% yield). $^1$H NMR (400 MHz, $CDCl_3$) δ 9.76 (q, *J* = 1.6 Hz, 1H), 4.12 (qd, *J* = 7.2, 1.2 Hz, 2H), 2.44 (tt, *J* = 7.3, 1.4 Hz, 2H), 2.30 (td, *J* = 7.4, 1.2 Hz, 2H), 1.65 (dtd, *J* = 8.6, 7.3, 5.8 Hz, 4H), 1.43 – 1.30 (m, 2H), 1.25 (td, *J* = 7.1, 1.2 Hz, 3H).

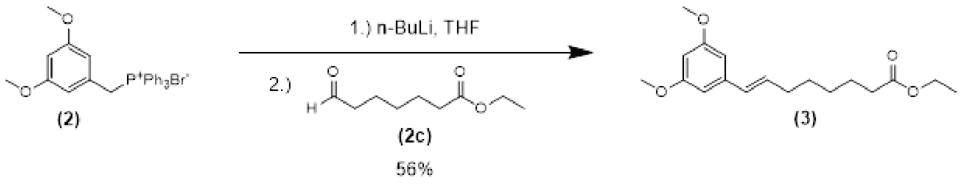


**Scheme S4. Synthesis of ethyl (E/Z)-8-(3,5-dimethoxyphenyl)oct-7-enoate (3)**Alkene (**3**) was synthesized as described by Sargent and Wangchareontrakul.[6] To a 100 mL oven dried two-neck flask under nitrogen atmosphere was added 5.00 g (10.1 mmol, 1 eq) phosphonium bromide **(2)** and 25 mL of dry THF. 6.2 mL (13 mmol, 1.3 eq) of 2.12 M n-BuLi in hexane was added dropwise and the solution was stirred at room temperature for 30 minutes. Then, 1.22 g ethyl 7-oxoheptanoate (7.07 mmol, 0.7 eq) was diluted in 12.5 mL of dry THF and added to the reaction dropwise. The reaction was stirred overnight before pouring the reaction mixture into ~150 mL of ice water. The mixture was extracted with 100 mL ethyl acetate three times. The organic layers were combined and washed once with brine then dried over $Na_2SO_4$. A rotary evaporator was used to remove solvent until approximately 5 mL of material remained. 10 mL of fresh ethyl acetate was added to the mixture and allowed to sit at room temperature overnight. The liquid was decanted from the white solids (TPPO) that precipitated. Fresh ethyl acetate was added to the mixture and allowed to sit for ~6 hours, and the liquid once again decanted. The decantates were pooled and the volatiles removed by rotary evaporation until ~8 mL of liquid remained. This was placed in a freezer (-20 °C) to precipitate more TPPO from which the liquid was decanted. This was repeated three more times before finally concentrating the product using a rotary evaporator. The crude product was purified by flash chromatography using a silica gel column and a solvent gradient of 0 – 10% ethyl acetate in hexanes. The product was concentrated under vacuum to yield 0.93 g (43%) of **(3)** as a mixture of E and Z isomers. MS: m/z = +329 (observed), [M+Na] = 329.4 (calculated). $^{1}$H NMR (400 MHz, $CDCl_3$) δ 6.49 (d, *J* = 2.3 Hz, 2H), 6.41 (d, *J* = 2.3 Hz, 1H), 6.33 (dt, *J* = 9.1, 2.3 Hz, 3H), 6.28 (s, 1H), 6.18 (dt, *J* = 15.8, 6.7 Hz, 1H), 5.62 (dt, *J* = 11.6, 7.2 Hz, 1H), 4.10 (qd, *J* = 7.2, 3.0 Hz, 3H), 3.76 (s, 10H), 2.37 – 2.23 (m, 5H), 2.19 (q, *J* = 7.1, 6.6 Hz, 2H), 1.69 – 1.56 (m, 4H), 1.51 – 1.31 (m, 8H), 1.23 (td, *J* = 7.1, 1.4 Hz, 4H), 0.92 (td, *J* = 7.4, 1.5 Hz, 1H). $^{13}$C NMR (101 MHz, $CDCl_3$) δ 174.03, 173.93, 173.88, 161.01, 160.64, 140.04, 139.76, 133.43, 131.50, 130.06, 129.07, 106.99, 106.90, 104.15, 99.33, 98.85, 64.28, 64.27, 60.33, 60.32, 55.43, 34.44, 32.85, 30.83, 29.74, 29.05, 28.99, 28.97, 28.83, 28.81, 28.71, 25.04, 25.00, 24.96, 19.28, 14.39, 13.84.

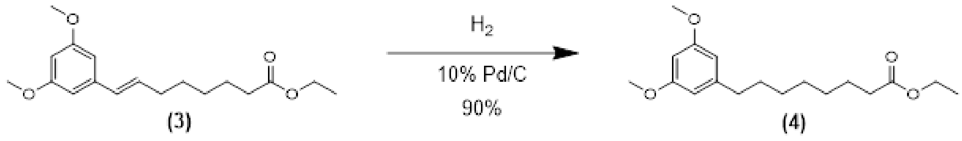


**Scheme S5. Synthesis of ethyl 8-(3,5-dimethoxyphenyl)octanoate (4)** Reduction of alkene (**3**) was achieved as described by Sargent and Wangchareontrakul.[6] 81 mg 10% Pd/C was added to an oven dried 100 mL flask and with a stir bar and the catalyst was wet with 1 mL of EtOAc. 0.69 g (2.26 mmol, 1 eq) of the alkene **(3)** was dissolved in 6 mL of EtOAc and 12 mL EtOH, then added to the flask while stirring. A balloon containing $H_2$ was attached to the top of the flask and the reaction was left stirring at room temperature for three days. Then, the contents of the flask were filtered through celite. The celite filter cake washed with EtOAc and EtOH. The combined filtrate and washes were concentrated using a rotary evaporator. The crude product was purified by flash chromatography using a silica gel column and eluent of 2 – 10% EtOAc in hexanes to afford 0.53 g (77%) of a clear oil. MS: m/z = +309 and +331 (observed), [M+H] = 309.4 (calculated), [M+Na] = 331.4 (calculated). $^{1}$H NMR (400 MHz, $CDCl_3$) δ 6.34 (d, *J* = 2.3 Hz, 2H), 6.29 (t, *J* = 2.3 Hz, 1H), 4.12 (q, *J* = 7.2 Hz, 2H), 3.78 (s, 6H), 2.54 (dd, *J* = 8.8, 6.7 Hz, 2H), 2.28 (t, *J* = 7.5 Hz, 2H), 1.60 (q, *J* = 7.4 Hz, 5H), 1.33 (h, *J* = 4.9, 4.1 Hz, 6H), 1.25 (t, *J* = 7.1 Hz, 3H). $^{13}$C NMR (101 MHz, $CDCl_3$) δ 174.02, 160.82, 145.41, 106.60, 97.71, 60.31, 55.37, 36.40, 34.51, 31.33, 29.27, 29.20, 25.10, 14.40.

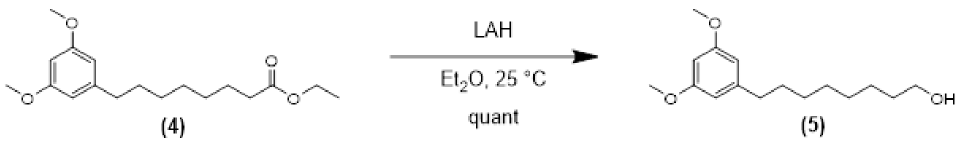


**Scheme S6. Synthesis of 8-(3,5-dimethoxyphenyl)octan-1-ol (5)** Reduction of ester (**4**) to the primary alcohol was achieved as described by Sargent and Wangchareontrakul.[6] To an oven-dried 50 mL round bottom two-neck flask with stir bar under an atmosphere of nitrogen was added 84 mg LAH (2.21 mmol, 1.3 eq). 7 mL of dry diethyl ether was slowly added to the flask while gently stirring. Next, 0.53 g (1.7 mmol, 1 eq) of ester was dissolved in 14 mL of dry ether and slowly added to the reaction flask. The mixture was stirred for 3 h at room temperature. After 3 h, the flask was placed on ice and quenched by the dropwise addition of saturated $NH_4Cl$. The reaction mixture was combined with an equal volume of saturated Rochelle salt solution and stirred overnight. The organic and aqueous phases were separated, and the aqueous phase extracted with 25 mL EtOAc three times. The organic layers were combined and washed with brine and dried over $Na_2SO_4$. Volatiles were removed by rotary evaporation and the product concentrated under vacuum to yield 0.43 g (95%) of material that was used in the next step without further purification. MS: m/z = +289 (observed), [M+Na] = 289.4 (calculated); m/z = +267 (observed), [M+H] = 267.4 (calculated). $^{1}$H NMR (400 MHz, $CDCl_3$) δ 6.34 (d, *J* = 2.3 Hz, 2H), 6.30 (t, *J* = 2.3 Hz, 1H), 3.78 (s, 6H), 3.63 (t, *J* = 6.6 Hz, 2H), 2.57 – 2.51 (m, 2H), 1.64 – 1.52 (m, 4H), 1.44 (s, 1H), 1.37 – 1.29 (m, 8H).

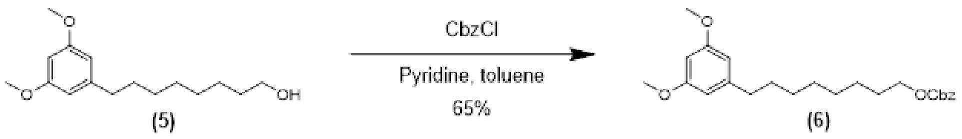


**Scheme S7. Synthesis of benzyl (8-(3,5-dimethoxyphenyl)octyl) carbonate (6)** Carboxybenzyl (Cbz)-protected intermediate (**6**) was synthesized according to the total synthesis of sorgoleone by Sargent and Wangchareontrakul.[6] To an oven-dried 25 mL round bottom flask and stir bar under nitrogen was added 0.43 g (1.6 mmol, 1 eq) of the octanol starting material **(5)** dissolved in 7.2 mL dry DCM. The solution was cooled in an ice bath and 0.39 mL (0.38 g, 4.8 mmol, 3 eq) of dry pyridine was added, followed by 0.30 mL (0.35 g, 2.1 mmol, 1.3 eq) of benzyl chloroformate. The reaction was left stirring overnight, warming to room temperature. The reaction mixture was diluted with EtOAc, washed with 5% citric acid, then washed with saturated $NaHCO_3$, and finally washed with brine before drying over $Na_2SO_4$. Volatiles were removed by rotary evaporation and the crude product purified by flash chromatography on a column of silica gel eluent 5 – 40% EtOAc in hexanes. Solvent was removed by rotary evaporation and the product concentrated under vacuum to give 0.42 g (65%) of **(6)** as a clear oil. 0.019 g of starting material was also recovered. $^{1}$H NMR (400 MHz, $CDCl_3$) δ 7.41 – 7.31 (m, 6H), 6.34 (d, *J* = 2.3 Hz, 2H), 6.29 (t, *J* = 2.3 Hz, 1H), 5.15 (s, 2H), 4.14 (t, *J* = 6.7 Hz, 2H), 3.78 (s, 6H), 2.53 (dd,

$J$ = 8.7, 6.7 Hz, 2H), 1.70 – 1.63 (m, 2H), 1.63 – 1.55 (m, 2H), 1.38 – 1.28 (m, 8H). $^{13}$C NMR (101 MHz, $CDCl_3$) δ 160.81, 155.39, 145.41, 135.48, 128.70, 128.61, 128.46, 106.59, 97.68, 69.58, 68.43, 55.35, 36.40, 31.34, 29.45, 29.32, 29.25, 28.76, 25.80.

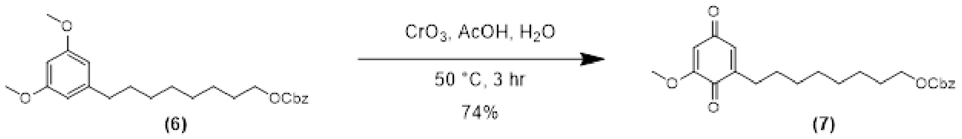


**Scheme S8. Synthesis of benzyl (8-(5-methoxy-3,6-dioxocyclohexa-1,4-dien-1-yl)octyl) carbonate (7)** Oxidation of intermediate (**6**) to benzoquinone (**7**) was adapted from the total synthesis of sorgoleone by Sargent and Wangchareontrakul.[6] 1.05 g (2.5 mmol, 1 eq) of the Cbz protected alcohol **(6)** was dissolved in 5 mL AcOH and added to a 50 mL flask in an atmosphere of nitrogen. In a separate vial, $CrO_3$ (1.06 g, 10.5 mmol, 4 eq) was dissolved in 2.5 mL acetic acid with 2.5 mL water, then added to the stirring reaction. The mixture was heated to 50 °C and covered with foil (to protect from light). After three hours, the heat was turned off and the reaction was allowed to cool to room temperature while stirring overnight. The reaction was diluted with ~40 mL of water and extracted with EtOAc three times. The organic layers were pooled, washed with saturated $NaHCO_3$, washed with water, and finally washed with brine before drying over $Na_2SO_4$. A rotary evaporator was used to remove the solvent, yielding the crude product as a dark yellow/brown oil. The crude product was purified by flash chromatography using a silica gel column using 10 – 50% EtOAc in hexanes as eluent. Solvent was removed by rotary evaporation and the material dried under vacuum giving 0.78 g (74%) of a yellow/orange oil that crystallized to a bright yellow solid upon standing. MS: m/z = +423 (observed), [M+Na] = 423.5 (calculated). $^1$H NMR (400 MHz, $CDCl_3$) δ 7.41 – 7.30 (m, 5H), 6.47 (dt, $J$ = 2.6, 1.5 Hz, 1H), 5.87 (d, $J$ = 2.4 Hz, 1H), 5.15 (s, 2H), 4.14 (t, $J$ = 6.7 Hz, 2H), 3.81 (s, 3H), 2.42 (ddd, $J$ = 8.8, 7.2, 1.5 Hz, 2H), 1.66 (t, $J$ = 7.2 Hz, 2H), 1.48 (q, $J$ = 7.4, 6.9 Hz, 2H), 1.39 – 1.27 (m, 8H). $^{13}$C NMR (101 MHz, $CDCl_3$) δ 187.80, 182.27, 158.99, 155.40, 147.63, 135.49, 133.05, 128.73, 128.64, 128.49, 107.26, 69.62, 68.40, 56.42, 29.28, 29.26, 29.18, 28.85, 28.76, 27.83, 25.78.

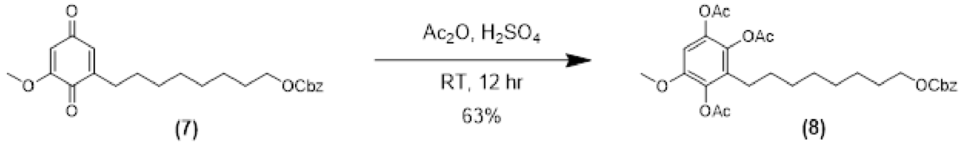


**Scheme S8. Synthesis of 3-(8-(((benzyloxy)carbonyl)oxy)octyl)-5-methoxybenzene-1,2,4-triyl triacetate (8)** Acetoxylation of benzoquinone (7) was performed as described by Sargent and Wangchareontrakul.[6] To an oven-dried 25 mL flask and stir bar under nitrogen was added 0.34 g (0.85 mmol, 1 eq) of the benzoquinone **(7)** dissolved in 4.9 mL (51 mmol, 60 eq) of $Ac_2O$. 69 µL (1.3 mmol, 1.5 eq) of concentrated $H_2SO_4$ was added dropwise while stirring. The reaction was protected from light and left stirring overnight at room temperature. Then, the reaction mixture was poured over ice (~100 mL) and the crude product extracted with EtOAc three times. The organic layers were pooled and washed with saturated $NaHCO_3$, washed with water, washed with brine, and finally dried over $Na_2SO_4$. Solvent was removed using a rotary evaporator. The reside was purified by flash chromatography using a silica gel column and a solvent gradient of 15 – 40% EtOAc in hexanes. Solvent was removed by rotary evaporation and the purified product dried under vacuum, yielding 0.29 g (63%) of the triacetate **(8)**; 75 mg of the quinone starting material was also recovered. MS: m/z = +567 (observed), [M+Na] = 567.6 (calculated). $^1$H NMR (400 MHz, $CDCl_3$) δ 7.40 – 7.30 (m, 5H), 6.70 (s, 1H), 5.15 (s, 2H), 4.13 (t, $J$ = 6.7

Hz, 2H), 3.78 (s, 3H), 2.45 – 2.37 (m, 2H), 2.32 – 2.24 (m, 9H), 1.68 – 1.61 (m, 2H), 1.46 – 1.39 (m, 2H), 1.37 – 1.23 (m, 10H). $^{13}$C NMR (101 MHz, $CDCl_3$) δ 168.48, 168.39, 168.23, 155.36, 149.29, 140.34, 136.07, 135.46, 133.97, 130.20, 128.69, 128.61, 128.44, 104.96, 69.57, 68.38, 56.34, 29.64, 29.20, 29.16, 29.05, 28.75, 25.76, 25.44, 20.86, 20.55, 20.40.

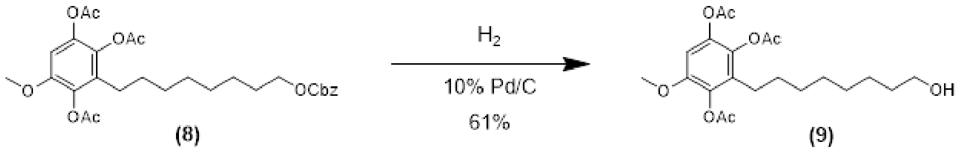


**Scheme S9. Synthesis of 3-(8-hydroxyoctyl)-5-methoxybenzene-1,2,4-triyl triacetate (9)** To an oven-dried 25 mL round bottom flask and stir bar was added 2 mg 10% Pd/C. The catalyst was wet with 0.5 mL of EtOAc. 30 mg (0.06 mmol, 1 eq) Cbz protected alcohol **(8)** was dissolved in 1 mL of EtOAc, then added to the flask while stirring. A balloon containing $H_2$ was attached to the top of the flask and the reaction was stirred at room temperature for four days, refilling the $H_2$ balloon as needed. After four days, the reaction mixture was diluted with EtOAc and the contents of the flask were filtered through celite. The celite filter cake was washed with EtOAc. The filtrate and washes were combined, and the solvent removed using a rotary evaporator. The crude product was purified by flash chromatography using a silica gel column and a solvent gradient of 10 – 60% EtOAc in hexanes. Solvent was removed by rotary evaporation and the purified product dried under vacuum to yield 23 mg (61%) of the alcohol **(9)**. MS: m/z = 433.4 (observed), [M+Na] = 433.5 (calculated). $^{1}$H NMR (400 MHz, $CDCl_3$) δ 6.70 (s, 1H), 3.78 (s, 3H), 3.63 (t, $J$ = 6.6 Hz, 2H), 2.45 – 2.37 (m, 2H), 2.30 (d, $J$ = 8.4 Hz, 6H), 2.26 (s, 3H), 1.55 (p, $J$ = 6.8 Hz, 2H), 1.48 – 1.39 (m, 3H), 1.29 (q, $J$ = 6.8, 5.9 Hz, 11H). $^{13}$C NMR (101 MHz, $CDCl_3$) δ 168.56, 168.47, 168.31, 149.32, 140.35, 136.09, 133.99, 130.24, 104.97, 63.10, 56.36, 32.87, 29.59, 29.30, 29.28, 29.03, 25.75, 25.46, 20.88, 20.57, 20.42.

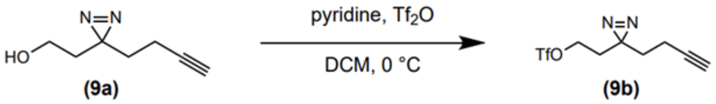


**Scheme S10. Synthesis of 2-(3-(but-3-yn-1-yl)-3H-diazirin-3-yl)ethyl trifluoromethanesulfonate (9b)** Triflated diazirine alkyne (**9b**) was synthesized according to a previously reported method.[7] An oven dried 25 mL flask and stir bar under an atmosphere of nitrogen was placed in an ice bath. 1.1 mL dry DCM was added to the flask and allowed to cool for 15 minutes. Then, 25 µL (27 mg, 0.19 mmol, 1.3 eq) of the alcohol linker 2-(3-but-3-ynyl-3H-diazirin-3-yl)-ethanol **(9a)** was added to the flask while stirring, followed by 16 µL (15 mg, 0.19 mmol, 1.3 eq) of dry pyridine, and finally 32 µL (54 mg, 0.19 mmol, 1.3 eq) triflic anhydride. The reaction was protected from light and stirred at 0 °C for 30 minutes. After 30 minutes, a red solid formed in the reaction mixture (triflate-pyridinium salt side product), and the reaction was quenched by dropwise addition of water. Stirring was stopped and the lower layer was removed using a syringe and concentrated using a rotary evaporator. The residue was suspended in pentane and filtered through $Na_2SO_4$ and the filtrate concentrated using a rotary evaporator. Remaining water was removed by azeotroping with anhydrous toluene, giving 0.036 g (69%) of the triflate **(9b)** which was used immediately in the next reaction without further purification or characterization.

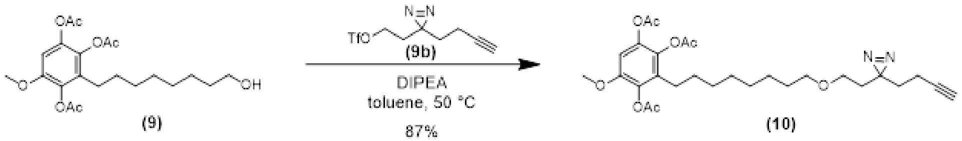


**Scheme S11. Synthesis of 3-(8-(2-(3-(but-3-yn-1-yl)-3H-diazirin-3-yl)ethoxy)octyl)-5-methoxybenzene-1,2,4-triyl triacetate (10)** To an oven-dried 50 mL flask and stir bar under nitrogen was added 58 mg (0.14 mmol, 1 eq) of the alcohol **(9)** dissolved in 0.5 mL dry toluene. The solution was cooled to 0 °C, then 0.14 mL (0.78 mmol, 5.5 eq) DIPEA was added to the flask. The freshly prepared triflate linker **(9b)** was dissolved in 0.5 mL dry toluene and added to the reaction flask and the reaction was protected from light. The ice bath was removed, and the reaction mixture was allowed to warm to room temperature. Once the reaction mixture reached room temperature, it was heated to 50 °C and left stirring at this temperature for 48 h. After 48 h, the reaction was allowed to cool to room temperature and diluted with 10 mL of water and 10 mL EtOAc. The layers were separated, and the aqueous layer extracted with EtOAc two times. The pooled organic layers were washed with citric acid two times, followed by saturated NaHCO3, and dried over $Na_2SO_4$. Solvent was removed by rotary evaporation. The crude product was purified by flash chromatography using a column of silica gel and a solvent gradient of 20 – 50% EtOAc in hexanes. Solvent was removed by rotary evaporation and the purified product dried under vacuum giving 0.065 g (87%) of **(10)** as a slight pale-yellow oil. MS: m/z = +553 and +525 (observed), [M+Na] = 553.6 (calculated), [M-$N_2$+Na] = 525 (calculated). $^1$H NMR (400 MHz, $CDCl_3$) δ 6.69 (s, 1H), 3.77 (s, 3H), 3.34 (t, $J$ = 6.6 Hz, 2H), 3.22 (t, $J$ = 6.3 Hz, 2H), 2.44 – 2.36 (m, 2H), 2.31 (s, 3H), 2.29 (s, 3H), 2.25 (s, 3H), 2.05 – 1.99 (m, 3H), 1.96 (t, $J$ = 2.6 Hz, 1H), 1.69 – 1.62 (m, 4H), 1.54 (t, $J$ = 7.1 Hz, 2H), 1.46 – 1.38 (m, 2H), 1.35 – 1.23 (m, 11H). $^{13}$C NMR (101 MHz, $CDCl_3$) δ 168.51, 168.42, 168.25, 149.31, 140.35, 136.08, 133.98, 130.28, 104.95, 83.02, 71.27, 69.12, 65.24, 56.35, 33.27, 32.88, 29.74, 29.44, 29.33, 29.10, 27.04, 26.25, 25.47, 20.88, 20.57, 20.42, 13.42.

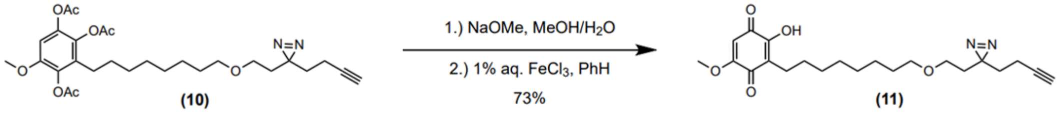


**Scheme S12. Synthesis of 3-(8-(2-(3-(but-3-yn-1-yl)-3H-diazirin-3-yl)ethoxy)octyl)-2-hydroxy-5-methoxycyclohexa-2,5-diene-1,4-dione (11)** 0.032 g (0.06 mmol, 1 eq) of the triacetate **(10)** was dissolved in 1 mL of dry MeOH and added to an oven-dried 25 mL flask with stir bar under an atmosphere of nitrogen. 43 µL (40 mg, 3.1 eq) NaOMe solution (25 wt% NaOMe in MeOH) was added to the flask while stirring. The reaction was protected from light and stirred at room temperature overnight. Then, the mixture was neutralized with Amberlite IR 120 resin. The solution was allowed to stir with the resin for 15 minutes before diluting with MeOH and filtered through a fritted disk. Solvents were removed solvent using a rotary evaporator. The residue was dissolved in 2 mL of benzene and 0.25 mL of 1% $FeCl_3$ aqueous solution was added, and the mixture was stirred vigorously at room temp for 3 hours. Next, the reaction mixture was diluted with water and the product extracted with EtOAc three times. The pooled organic layers were washed with brine and dried over $Na_2$SO4 before evaporating the solvent using a rotary evaporator. The residue was purified by flash chromatography using a silica gel column and solvent gradient of 0 – 10% MeOH in DCM. Fractions containing the product were concentrated using a rotary evaporator and dried under vacuum to give 0.018 g (73%) of semi-pure material. The semi-pure material was dissolved in 300 µL of DMSO for purification by HPLC (Agilent) as follows: 100 µL of the DMSO solution was injected for purification using a semi-prep C18 column with a solvent gradient of 5 – 95% MeCN with 0.1% TFA in $H_2O$ with 0.1% TFA over 60 minutes. Fractions containing the pure product were pooled, the organic solvent removed by rotary evaporation, and lyophilized to yield 3 mg of the final product **(11)**. MS: m/z = -401 (observed), [M-H] = 401.5 (calculated). $^1H$ NMR (400 MHz, $CDCl_3$) δ 7.21 (s, 1H), 5.84 (s, 1H), 3.86 (s, 3H), 3.35 (t, *J* = 6.6 Hz, 2H), 3.23 (t, *J* = 6.4 Hz, 2H), 2.47 – 2.41 (m, 2H), 2.03 (td, *J* = 7.6, 2.6 Hz, 2H), 1.97 (t, *J* = 2.7 Hz, 1H), 1.70 – 1.64 (m, 4H), 1.58 – 1.51 (m, 2H), 1.45 (t, *J* = 7.5 Hz, 2H), 1.37 – 1.24 (m, 9H). $^{13}C$ NMR (101 MHz, $CDCl_3$) δ 161.14, 151.53, 119.22, 102.18, 71.20, 68.99, 65.13, 56.79, 33.17, 32.76, 29.72, 29.60, 29.46, 29.33, 29.30, 28.00, 26.09, 22.60, 13.31.

### *Supplemental Figures*

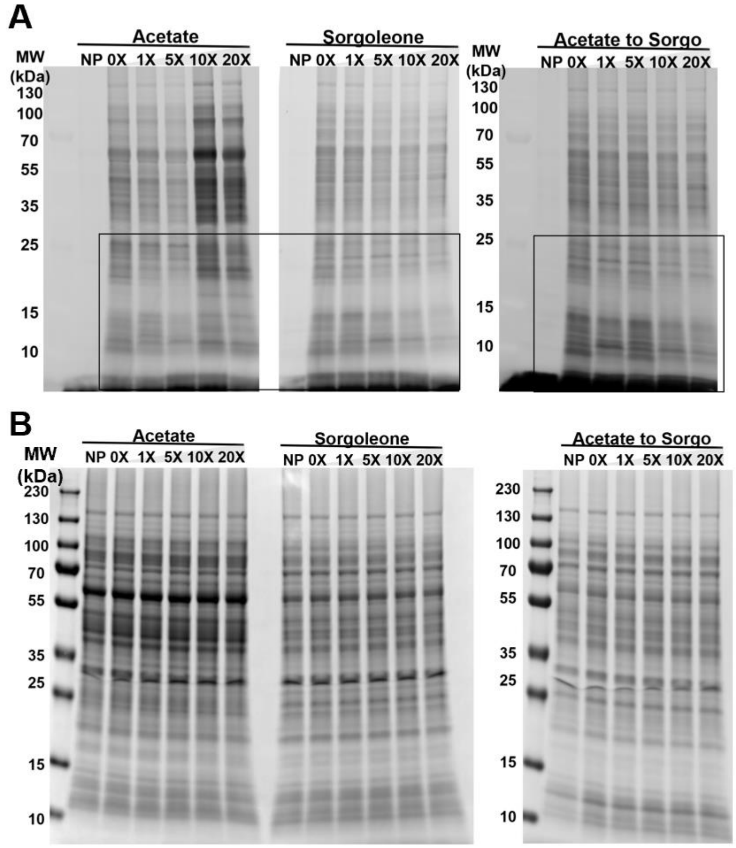


**Figure S1.** A. In-gel fluorescence analysis of SO-1clarified lysate grown on either acetate (“Acetate”), sorgoleone (“Sorgoleone”) or started with acetate and switched to sorgoleone (“Acetate to Sorgo”). Lysates sorgoleone treated with either DMSO (-) or the maximum concentration of probe competitor sorgoleone (+, 20x the concentration of the probe, 500 µM) followed by probe labelling. Labels above the lanes indicate the carbon source with which SO-1 cells were originally cultured (“Acetate” – acetate (20 mM), “Sorgo – sorgoleone (2 mM)”, “AtoS” – cultured initially with acetate (20 mM) then sorgoleone (1 mM)). Arrows indicate loss of fluorescent signals in competed lanes. B In-gel fluorescence analysis of SO-1 clarified lysate with either DMSO (0X) or a range of probe competitor sorgoleone (0X, 1X, 5X, 10X, 20X correspond to 0, 25, 125, 250, 500 µM sorgoleone respectively) followed by probe labelling. B. Gels used in fluorescent analysis with Coomassie stain to serve as a protein loading control.

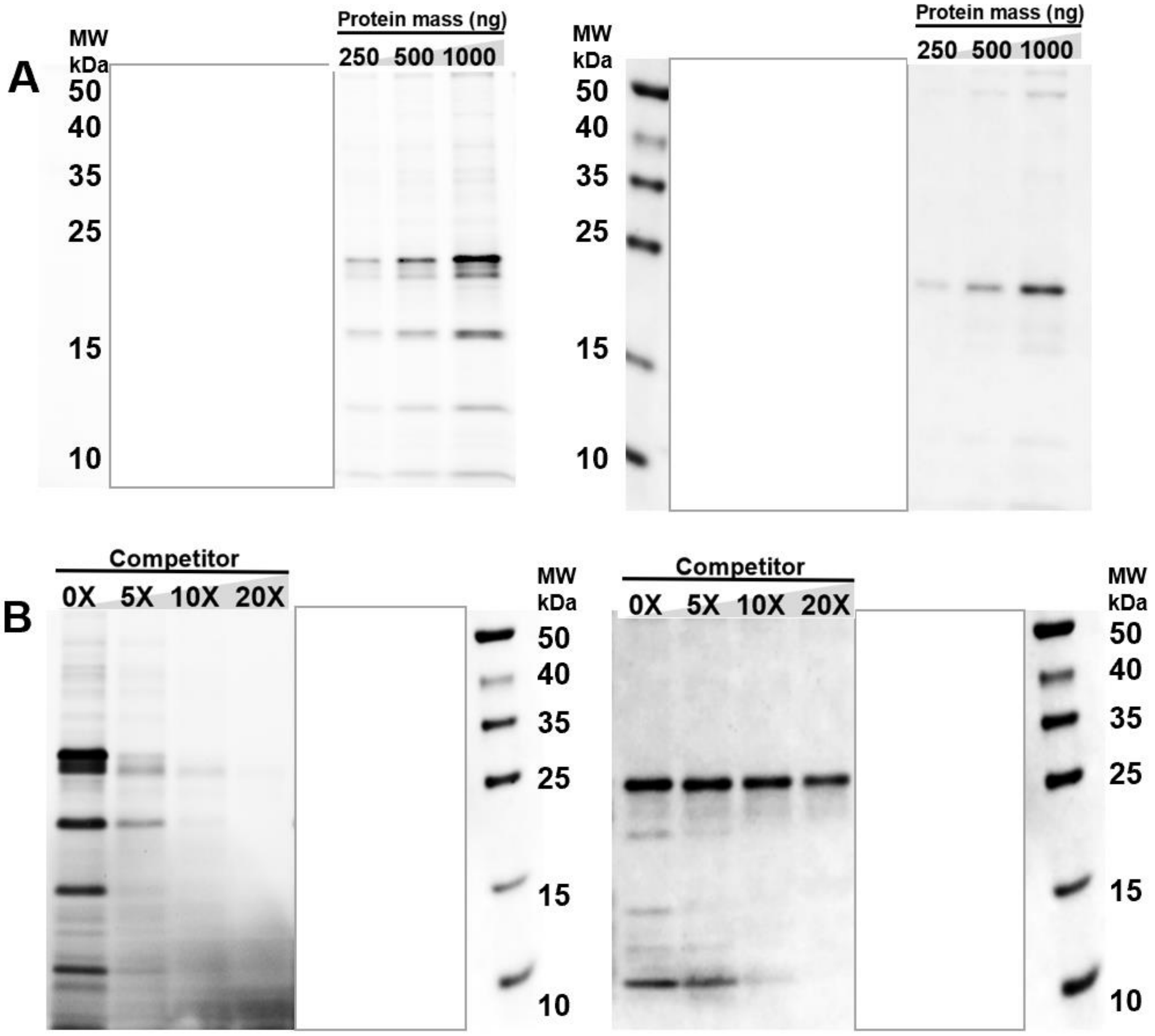


**Figure S2.** Whole gel images of the competition analysis shown in Figure 6B. A. In vitro protein labeling of recombinantly-expressed OH685_09420 (right panel is Cy3 fluorescence and left panel is the Coomassie stained gel) B. Competition assay (left) of recombinantly-expressed OH685_09420 labeling (right panel is Cy3 fluorescence and left panel is the Coomassie stained gel)

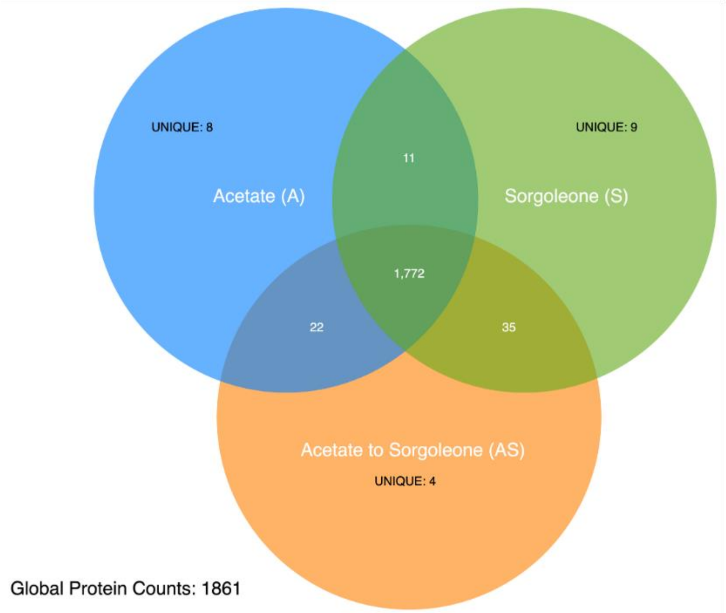


**Figure S3**. Venn diagram of shared and unique proteins identified through global proteomic analysis of *A. pittii* cultured on different carbon sources.

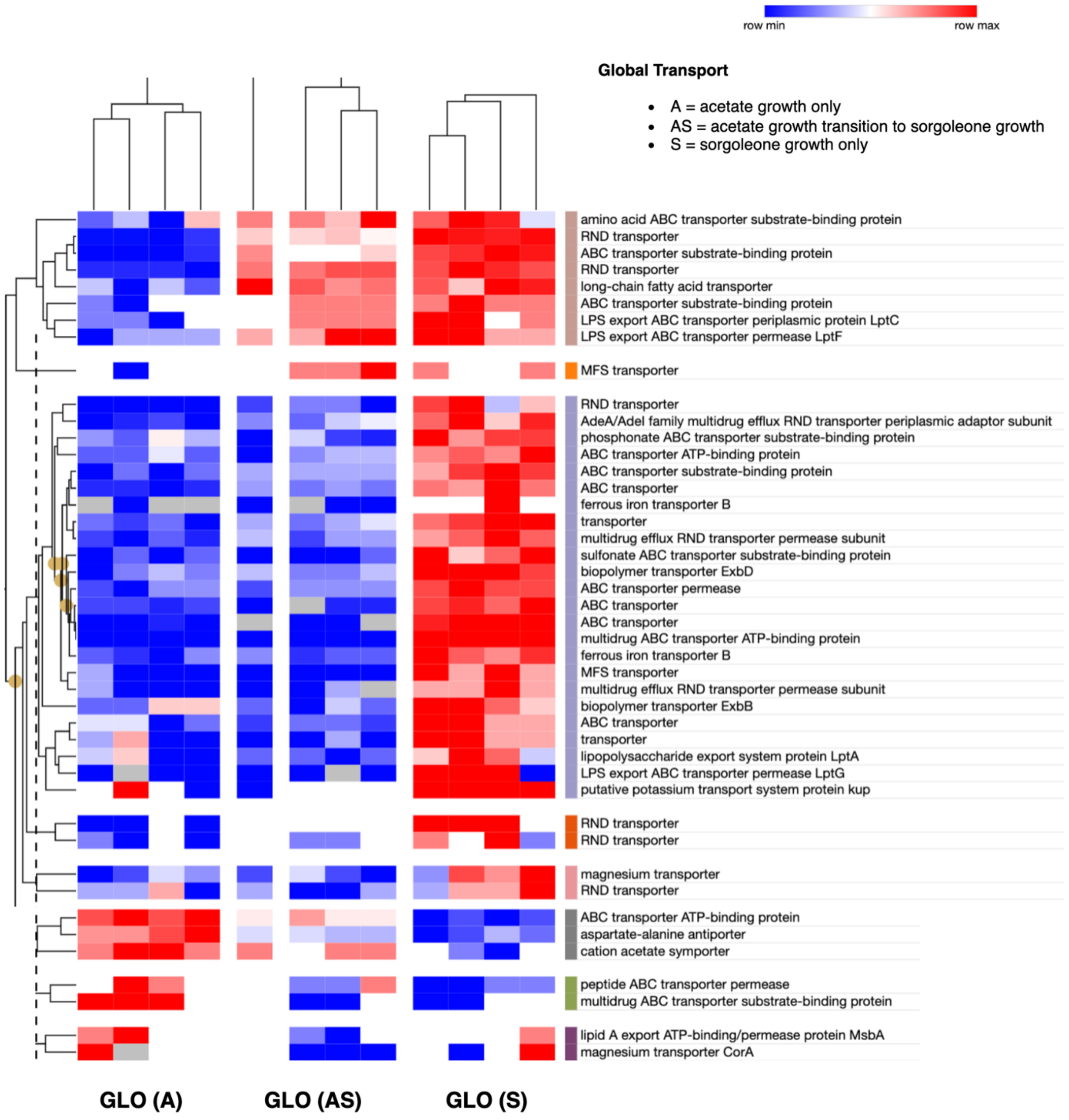


**Figure S4**. Global proteomic transporter profile on variable carbon transition growth medium.

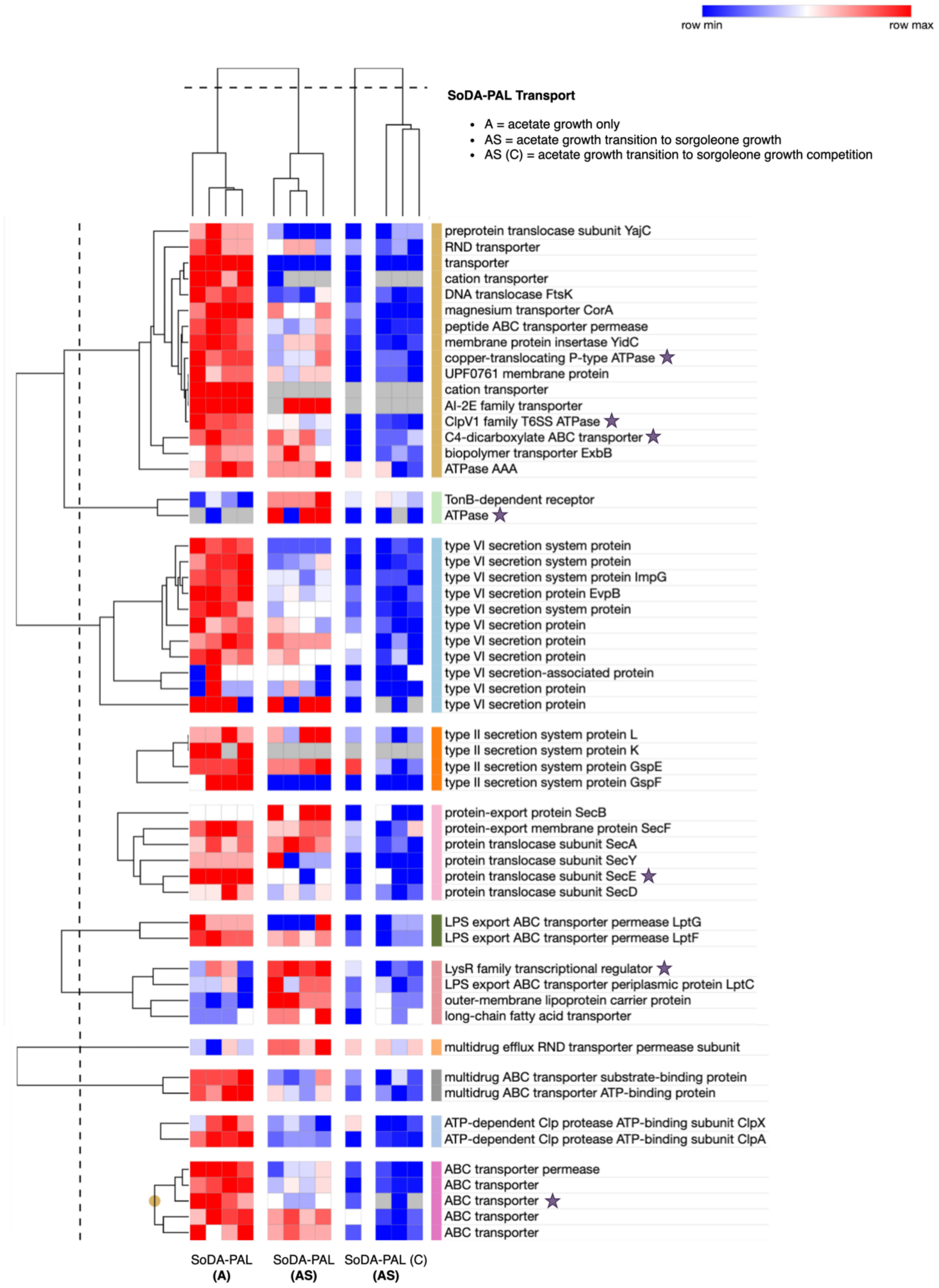


**Figure S5**. SoDA-PAL proteomic transporter profile on variable carbon transition growth medium. Stars indicate specific targets from our list of 137 significantly competed proteins.

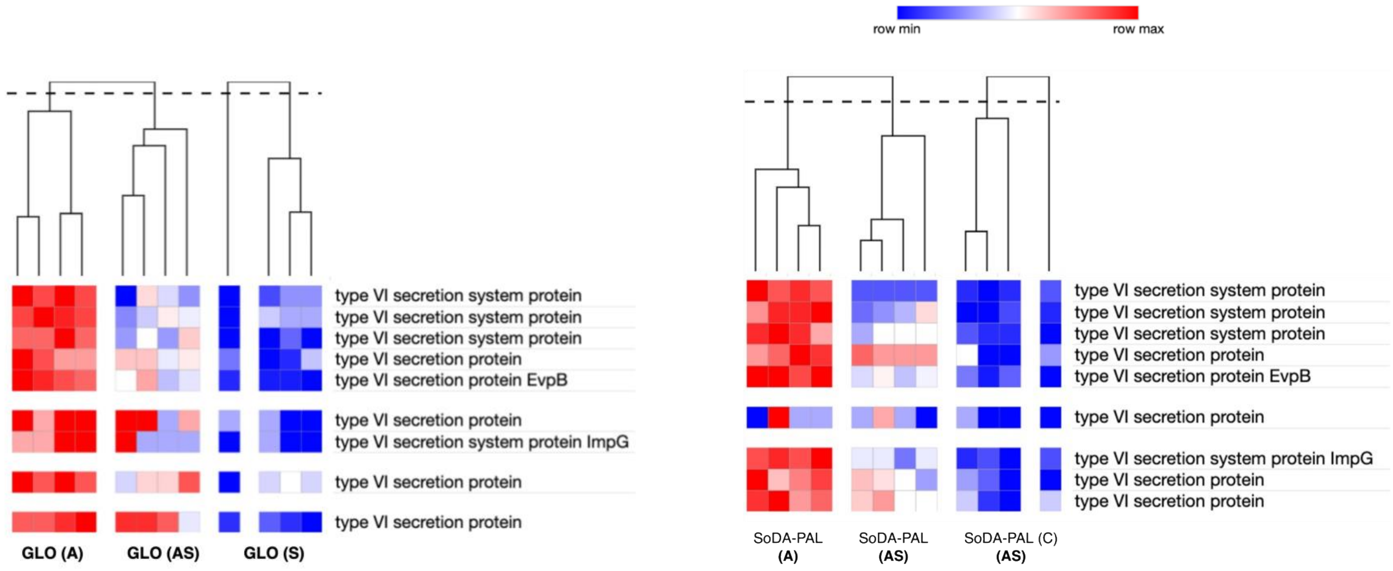


**Figure S6**. Global vs SoDA-PAL relative expression profile of type VI secretion system transport coverage

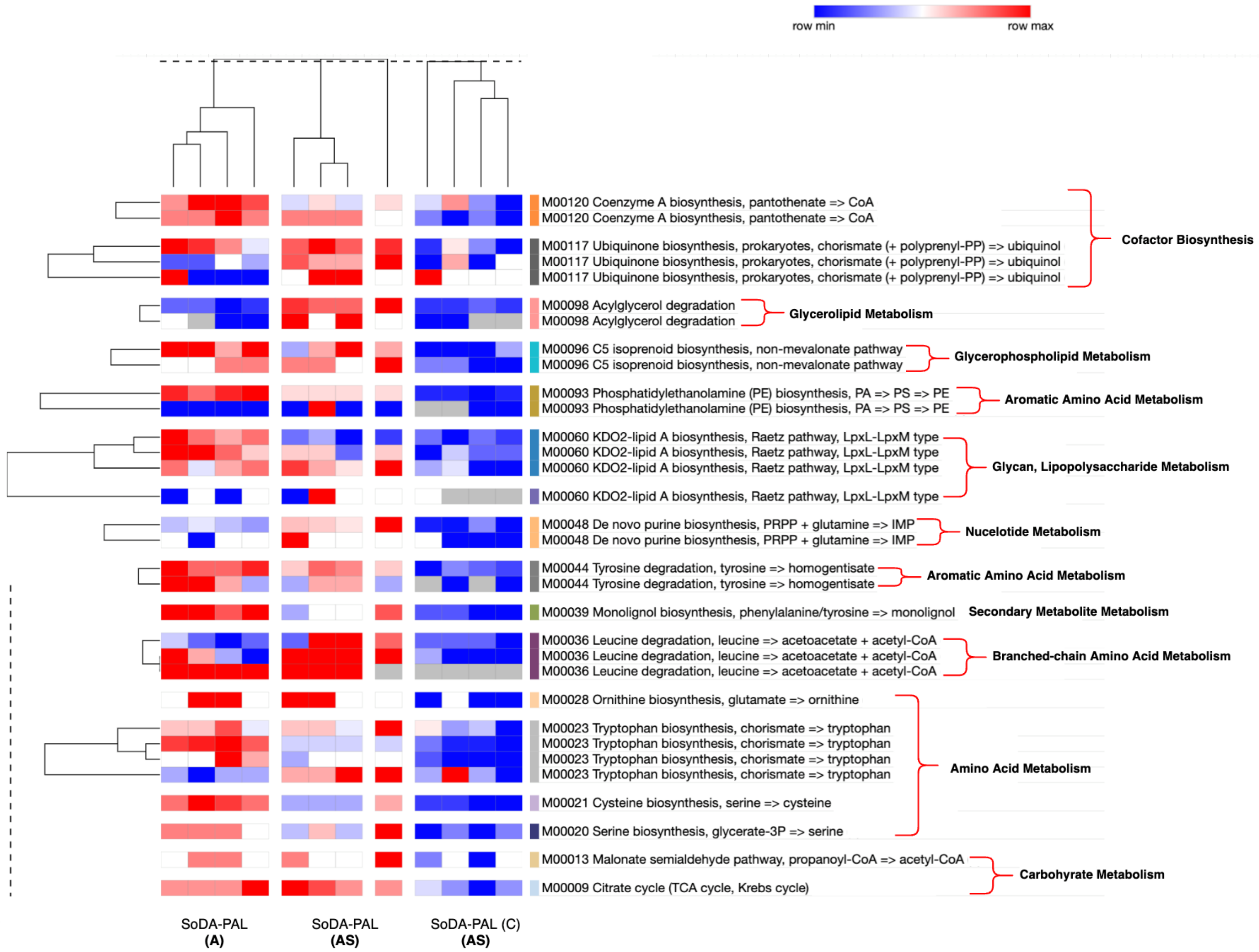


**Figure S7**. Proteins with secondary metabolism and lipid metabolism functions identified by SoDA-PAL in proteomes from *A. pittii* cultured under acetate only (A) and acetate to sorgoleone (AS) growth conditions.

**References:**

(1) Zarrabi, S.; Mahmoodi, N. O.; Marvi, O. Transesterification via Baeyer–Villiger oxidation utilizing potassium peroxydisulfate (K2S2O8) in acidic media. *Monatshefte für Chemie - Chemical Monthly* **2010**, *141* (8), 889-891. DOI: 10.1007/s00706-010-0338-9.

(2) Bosone, E.; Farina, P.; Guazzi, G.; Innocenti, S.; Marotta, V.; Valcavi, U. New Synthesis of Methyl 7-Oxoheptanoate: An Useful Intermediate for the Preparation of 2-(6-Methoxycarbonylhexyl)-cyclopent-2-en-1-one. *Synthesis* **1983**, *1983* (11), 942-944. DOI: 10.1055/s-1983-30584.

(3) Wong, M. L. J.; Sterling, A. J.; Mousseau, J. J.; Duarte, F.; Anderson, E. A. Direct catalytic asymmetric synthesis of alpha-chiral bicyclo[1.1.1]pentanes. *Nat Commun* **2021**, *12* (1), 1644. DOI: 10.1038/s41467-021-21936-4 From NLM Medline.

(4) Matsuura, B. S.; Keylor, M. H.; Li, B.; Lin, Y.; Allison, S.; Pratt, D. A.; Stephenson, C. R. A scalable biomimetic synthesis of resveratrol dimers and systematic evaluation of their antioxidant activities. *Angew Chem Int Ed Engl* **2015**, *54* (12), 3754-3757. DOI: 10.1002/anie.201409773 From NLM Medline.

(5) Schafroth, M. A.; Rummelt, S. M.; Sarlah, D.; Carreira, E. M. Enantioselective Iridium-Catalyzed Allylic Cyclizations. *Org Lett* **2017**, *19* (12), 3235-3238. DOI: 10.1021/acs.orglett.7b01346 From NLM PubMed-not-MEDLINE.

(6) Sargent, M. V.; Wangchareontrakul, S. The synthesis of the first natural host germination stimulant for Striga asiatica(witchweed). *Journal of the Chemical Society, Perkin Transactions 1* **1990**, (5). DOI: 10.1039/p19900001429.

(7) Flaxman, H. A.; Chang, C. F.; Wu, H. Y.; Nakamoto, C. H.; Woo, C. M. A Binding Site Hotspot Map of the FKBP12-Rapamycin-FRB Ternary Complex by Photoaffinity Labeling and Mass Spectrometry-Based Proteomics. *J Am Chem Soc* **2019**, *141* (30), 11759-11764. DOI: 10.1021/jacs.9b03764 From NLM Medline.
